# A generalizable normative deep autoencoder for brain morphological anomaly detection: application to the multi-site StratiBip dataset on bipolar disorder in an external validation framework

**DOI:** 10.1101/2024.09.04.611239

**Authors:** Inês Won Sampaio, Emma Tassi, Marcella Bellani, Francesco Benedetti, Igor Nenadic, Mary Phillips, Fabrizio Piras, Lakshmi Yatham, Anna Maria Bianchi, Paolo Brambilla, Eleonora Maggioni

**Affiliations:** Department of Electronics, Information and Bioengineering, Politecnico di Milano, Milan, Italy; Department of Neurosciences and Mental Health, Fondazione IRCCS Ca’ Granda Ospedale Maggiore Policlinico, Milan, Italy; Department of Neurosciences, Biomedicine and Movement Sciences, Section of Psychiatry, University of Verona, Verona, Italy; Division of Neuroscience, Unit of Psychiatry and Clinical Psychobiology, IRCCS Ospedale San Raffaele, Milan, Italy; Department of Psychiatry and Psychotherapy, Philipps-University Marburg, Marburg, Germany; Department of Psychiatry, University of Pittsburgh School of Medicine, Pittsburgh, PA, USA; Fondazione IRCCS Santa Lucia, Roma, Italy; Department of Psychiatry, University of British Columbia, Vancouver, BC, Canada; Department of Pathophysiology and Transplantation, University of Milan, Milan, Italy

**Author notes:** Equally contributing (Paolo Brambilla, Eleonora Maggioni).

**Keywords:** Normative Modelling, Anomaly Detection, Multi-site Harmonization, Psychiatric Disorders, Brain MRI

## Abstract

The heterogeneity of psychiatric disorders makes researching disorder-specific neurobiological markers an ill-posed problem. Here, we face the need for disease stratification models by presenting a generalizable multivariate normative modelling framework for characterizing brain morphology, applied to bipolar disorder (BD). We employed deep autoencoders in an anomaly detection framework, combined with a confounder removal step integrating training and external validation.

The model was trained with healthy control (HC) data from the human connectome project and applied to multi-site external data of HC and BD individuals. We found that brain deviating scores were greater, more heterogeneous, and with increased extreme values in the BD group, with volumes prominently from the basal ganglia, hippocampus and adjacent regions emerging as significantly deviating. Similarly, individual brain deviating maps based on modified z scores expressed higher abnormalities occurrences, but their overall spatial overlap was lower compared to HCs.

Our generalizable framework enabled the identification of subject- and group-level brain normative-deviating patterns, a step forward towards the development of more effective and personalized clinical decision support systems and patient stratification in psychiatry.

## 1. Introduction

Psychiatric disorders, as described in the current categorical classification system, are highly heterogeneous marked by a complex interplay of genetic and environmental factors that lead to altered physiological mechanisms [1]–[3]. Many neuroimaging studies have attempted to objectively characterize these disorders by searching for brain markers that could support diagnosis or disease management [4]–[10]. Nonetheless, no clinically useful markers have emerged until now [11]. For instance, brain models of bipolar disorder (BD) are currently being investigated, but the overall findings appear fragmented [12], [13]. A recurrent problem lies in the inability to generalize findings within a patient population, as group-level diagnostic effects have been shown to not replicate at subject level [14] and appear to be shared between different diagnostic groups [15]–[17]. Thus, delineating disorder-specific neurobiological patterns is challenging as the categorization of psychiatric disorders into well-defined diagnostic groups was not guided by neurobiological evidence [18], [19]. Accordingly, the study of brain morphological markers of psychiatric disorders should account for the uncertainty associated with the diagnostic labels and move away from classic case-control group comparisons to personalized normative-based statistical inferences [20]–[22].

Deep learning (DL) autoencoder (AE) models are widely employed in anomaly detection frameworks and have emerged has suitable multivariate models for brain normative frameworks [23]–[25]. AEs are encoder-decoder models based on artificial neural networks designed to capture relevant regularities in data through the minimization of a reconstruction error (RE). The REs are fully traceable, enabling the identification of specific brain regions with higher deviations from the norm. Thus, leveraging this normative-based anomaly detection approach effectively attenuates the lack of interpretability associated with their “black box” nature. In addition, model interpretability can be further increased through the application of the AE on confounder-free data [26].

By leveraging these promising modelling tools and a large multi-site T1-weighted structural magnetic resonance imaging (sMRI) data of healthy controls (HC) and individuals with BD, our study proposes a robust and innovative personalized medicine framework for improving the complex clinical management of BD (and other mental disorders). A shift from a disease-centred to a patient-centred paradigm is promoted via the development of a generalizable, and extendable AE-based brain normative modelling and anomaly detection framework. For the first time, the proposed data processing pipeline includes a confounders’ removal step fully generalizable to external datasets and the normative model integrates cortical thickness (CT), gray matter (GMV), and white matter volumes (WMV) features.

We developed an end-to-end pipeline to manage both biological and site-related confounding sources embedded in the external validation (EV) framework [27], allowing its application to new external data. The normative model framework was trained on region-based brain morphological features to integrate CT, GMV and WMV, from the human connectome project young adults (HCP-YA) data [28] and applied to multi-site data from the StratiBip network [16], including HC and subjects with BD. Subject-level REs were extracted and compared between HC and BD to assess and characterize deviating brain patterns in affected individuals, under the hypothesis that individuals affected with BD would deviate more than HCs from the HCP-YA normative population in some of the brain features. We hypothesized that the AE-based normative model built on HCP-YA data would be a robust and effective tool to identify and characterize subject-level and group-level patterns of brain alterations.

## 2. Related Work

Numerous techniques have been proposed for brain normative modelling and anomaly detection, which we distinguish here as regression-based and DL-based. Unsupervised deep learning models for anomaly detection are mostly based on AEs or Generative Adversarial Networks [25]. According to a recent review, most DL-based anomaly detection techniques for brain medical imaging have been developed for lesion and tumor detection or for brain segmentation, taking raw images and volumes as input [29]. Few examples can be found in literature applying this framework to psychiatric disorders, where brain alterations are subtle and not explicitly present. The first to develop such an application with deep AEs was *Pinaya et al*. [23], training an AE model with brain morphological features from healthy controls and then employing an anomaly detection framework to study brain normative deviations of schizophrenic and autistic patients. In the same line, based on an adversarial AE model, the same author studied brain morphological deviations from patients with Alzheimer’s disease and mild cognitive impairment [24]. More recently, a basic autoencoder was employed as a normative data-driven feature learner and applied to extract data-driven brain-deviating scores [30]. In the latter work, the AE was trained with brain volumetric features from healthy controls and then the test set reconstruction errors associated with controls and subjects affected by bipolar disorder were extracted and fed to a feature selection module and a random forest classifier. Besides the described studies, most normative modelling approaches developed for psychiatric disorders have applied regression methods. In this case, normative brain curves have been mapped mainly using Gaussian process regression (GPR), first proposed for normative modelling in [31], and since then has been extensively used [14], [20]. Differently, *C. J. Fraza et al*. [32] proposed warped Bayesian linear regression as an improvement upon the latter GPR, which was successfully implemented in the work developed by *S. Rutherford et al*. [22]. Other regression-based methods have also been proposed, such as generalized additive models [33], [34]; nevertheless, all these methods are univariate, since they fit a separate regression line to each brain region and therefore do not address the interdependences among brain regions [35]. Conversely, multivariate approaches can overcome this issue by facilitating the study of pattern-wise brain changes [36]. In *R. Ge et al*. [37], a comparative analysis of eight algorithms, including the aforementioned methods, identified multivariate fractional polynomials (MFP) as the most effective model; still, deep learning models surpass MFPs in capacity and in handling highly complex multivariate relationships.

In summary, the majority of anomaly detection techniques developed for studying psychiatric disorders have relied on regression methods, which are limited in their capacity to model complex multivariate relationships. Few studies have employed DL-based techniques to investigate brain morphological anomalies, and those that have failed to address a critical challenge in psychiatry: the heterogeneity of diagnostic groups. The present study aims to fill this gap by proposing an end-to-end normative framework based on deep-AEs and statistical inference methods to study both within-group heterogeneity and between-group discrimination.

## 2. Materials and methods

The data analysis workflow is schematized in Fig. 1. We extracted brain regional features from sMRI data that were fed into an embedded confounder removal (CR) pipeline that integrated training with the external test set, comprising both biological and site confounding effects removal. Then the normative AE model was trained with the confounder-free HCP-YA training set features. The StratiBip test set REs were extracted from the normative model and both subject’s and brain features-by-group mean deviation scores (MDS) were calculated employing the *mean square error*. In the group-level analysis, we evaluated the MDS group’s discriminative power, identified significant deviating neuroanatomical patterns in the BD group, and characterized both groups in terms of RE heterogeneity and extreme deviating values. Then, we built personalized subject-level brain deviating maps for all test subjects via modified z scores (mZ) transformation and studied individual abnormalities and groups’ spatial maps overlap.

**Fig 1.**
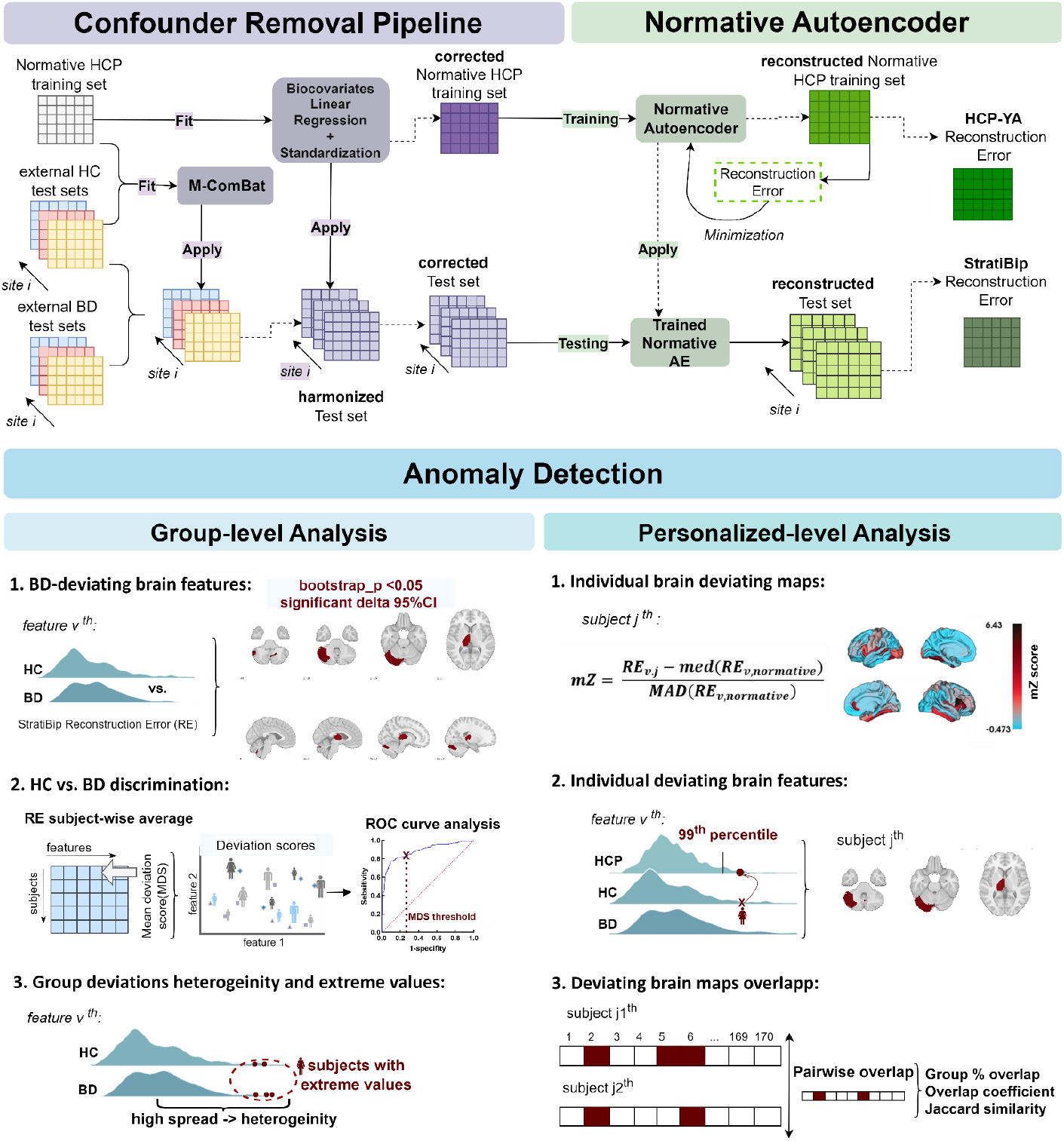
Normative modelling framework: data confounders’ removal and normative anomaly detection pipeline.

### 2.1 Datasets

#### Normative Training set: HCP dataset

Our training set was obtained from the Human Connectome Project Young Adults (HCP-YA) public dataset, *1200 Subjects Data Release (S12000 Release,March 2017)* [28], available on the connectomeDB platform (*https://db.humanconnectome.org*) [38]. The retrieved data consisted of 3T T1-weighted sMRI scans from 1109 HC subjects aged between 22 and 37 years. For this dataset, we obtained the *Restricted Data Access Authorization* by signing and agreeing to the WU-Minn HCP Terms. All methods developed and publication of source codes comply with the obligations and regulations of those terms.

#### StratiBip Dataset: External Test set

The external test set consisted of data collected as part of the StratiBip network, an initiative promoted by PB and EM that originated from the ENPACT network [16]. The StratiBip dataset results from the post-hoc integration of multi-site clinical and neuroimaging data collected from HC and subjects affected with BD, more details can be found in supplementary information.

The sMRI data used as external test set was acquired from 550 subjects (363 HC and 187 with BD) across 7 sites using T1-weighted sequences on 3T MRI scanners. Each site employed its own resources, protocols, and sequences (Table S12). Consistent with the HCP training sample, only young adults were included, from 22 to 37 years old (Table S1-S2).

### 2.2 sMRI pre-processing

All sMRI data were pre-processed in Matlab R2018a (The Mathworks, Inc®) environment. Firstly, T1-weighted images underwent a visual quality check and were converted from DICOM to NIFTI format. Following, the pre-processing was performed using the statistical parametric mapping software (SPM12) version 7771 [39], available at (http://www.fil.ion.ucl.ac.uk/spm/software/spm12/), and the computational anatomy toolbox add-on (CAT12) version 12.7 [40]. The detailed pre-processing pipeline is described in Supplementary Information. The pre-processed volume-based images were used to extract global measures as total intracranial volumes (TIV), regional cortical thickness measures for the Desikan-Killiany-Tourville (DK40) cortical atlas map [41], consisting of 68 ROIs (Table S13) and regional tissue volumes for the CoBra volume atlas map [42], provided by the Computational Brain Anatomy Laboratory at the Douglas Institute (CoBra Lab). The inclusion of volumetric measures was based entirely on the fully automated CAT12 processing pipeline. Therefore, all CAT12 volumetric estimations (WMV and GMV) using the CoBra atlas were included without any selection based on prior knowledge. CAT12 estimates WMV for GM regions and vice versa, using subject-specific tissue probability maps. WMV estimates for GM regions were interpreted as volume estimations for WM areas adjacent to the specific regions, and vice versa. A total of 50 GMV estimations (Table S14) and 52 WMV estimations (Table S15) were considered. The resulting GMV, WMV, and CT features were subject to the following processing steps.

### 2.3 Confounder Removal Pipeline

#### 2.3.1 Multi-site M-ComBat Harmonization

In the present work, we developed a framework for the harmonization of external test sets, i.e., with data collected in sites different from the normative training set. Site effects represent latent encoded information within MRI data associated with inter-site differences in MRI scanners and acquisition protocols. These differences make data not directly comparable, mask the biological effects of interest, e.g., diagnosis, and, most importantly, are learnable confounding features that significantly affect ML models and analysis [43]. In our study, the harmonization pipeline was developed to harmonize GM and WM volumes and CT features from the multi-site StratiBip external test set with the HCP-YA training set. This step was aimed to remove both intra-test set and inter-set differences, unlocking the possibility of applying the trained AE normative model in an external validation framework and performing reliable subject- and group-level statistical inferences. The pipeline was based on the ComBat (Combatting Batch Effects) tool, described below.

##### ComBat model

ComBat [44] is a harmonization method widely employed for neuroimaging datasets and particularly robust for small sample sizes [45], [46]. It uses an empirical bayes (EB) framework to estimate model parameters for each included site, assuming both additive and multiplicative site effects on data, *γ*_*iv*_, *δ*_*iv*_, for the *i*^*th*^site, *j*^*th*^ subject, and *v*^*th*^ feature *y* :

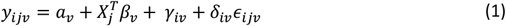

Furthermore, it allows for the preservation of subject-specific biological covariates, *X*_*j*_. The two site effect parameters are estimated from the standardized biocovariates-free data and then used to adjust the original data, as shown in Eq. 2:

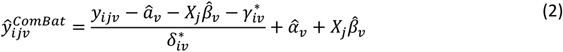

In the original ComBat model, the adjusted data is transformed to a location and scale related to the overall mean and pooled variance of the estimated site effects. Hence, to harmonize data, ComBat depends on the sites available at the moment of estimation, enabling its application exclusively in internal validation frameworks [47]–[49]. This issue is overcome in M-ComBat which gives the possibility to shift samples to a pre-determined reference batch location, 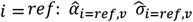, which we have employed for ML-EV frameworks as done before in [50].

##### Harmonization pipeline

The proposed harmonization pipeline is shown in Fig. 1. As we work in a normative modelling context, the StratiBip site-effect estimation was performed exclusively on the HC portion of the StratiBip test set (N=363), *y*_*ij=HC,v*_, using the HCP-YA normative training set as the reference *i = HCP*. In the site-effect estimation stage, the model starts by standardizing data with the HCP-YA statistics, 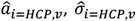, while accounting for biological covariates at net of site for all included subjects 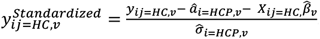. Next, additive and multiplicative site effects were estimated using the EB framework and then applied in the correction stage to harmonize the StratiBip external test set (relative to both HC and BD). The harmonization of the test set was performed as indicated in Eq. 2, using the feature mean, standard deviation, and biocovariates coefficients computed on the HCP-YA reference set.

The python-based *neurocombat* functions made available in (https://github.com/Jfortin1/neuroCombat) by F.P. Fortin were adapted and integrated into a python classe available in (https://github.com/inesws/neurocombat_pyClasse), denominated *neurocombat_pyClasse*, compatible with *sklearn Pipelines* and with *fit(), transform()* methods for its application in CV frameworks.

##### Feature harmonization

Using the pipeline described above, we harmonized TIV, WMV, GMV, and CT features of the StratiBip test set with the reference HCP-YA training set. For all feature sets, age and sex were included as biocovariates to preserve, while for volume features, previously harmonized TIV was also included. First, raw TIV measures were harmonized together with other extracted global measures. Then, regional volumes and CT features were separately harmonized. More detailed information is available in Supplementary Information.

##### Harmonization pipeline validation

To ascertain the harmonization success, we proceed with a series of validation analyses. The compliance with the following criteria was assessed: 1) successful and efficacy of site effects removal, 2) total preservation of biological covariates after M-ComBat harmonization. To evaluate 1) we checked if site differences and effects identified before data harmonization were effectively removed after M-ComBat application. We employed Kruskal Wallis ANOVA tests to compare mean feature-type distributions among sites and a site classification paradigm with a support vector machine learning model, before and after harmonization. To evaluate 2) we study the significance of age, sex, and diagnosis effects on raw and harmonized data with linear regression models to assert their stability after M-ComBat harmonization. A more detailed description of this and complementary analyses is available in Supplementary Information.

#### 2.3.2 Biological covariates removal via linear regression

After data harmonization, we proceeded with the removal of variance associated with age and sex biocovariates from regional volume and CT features, and harmonized TIV from volume features, via standard LR [51], [52]. We considered the outlined biological covariates as confounding variables as these are implicitly encoded in neuroimaging data and would contribute to a source ambiguity problem of the later developed AE model. We embedded the LR estimations and corrections in the EV, consistently with the proposed CR pipeline. The LR coefficient estimations were performed exclusively on the HCP-YA training set, and the estimated effects were removed from both HCP-YA training set and StratiBip test set [53], [54]. After this step, data is referred to as *corrected*. More detailed information can be found in Supplementary Information.

### 2.4 Autoencoder Normative Model

After data has been adjusted for the identified confounders, the following step is the implementation of the normative AE-based model.

#### AE for normative modelling

The implementation of AEs for normative modelling is within the scope of methods for normality feature learning by characterizing regular feature patterns [25]. An AE has an encoder-decoder architecture based on artificial neural networks and is widely used for data embedding representation learning. In the normative framework, the model is trained in an unsupervised fashion to learn to represent normative data by optimizing a generic objective function that minimizes the model reconstruction error. Then, by employing an anomaly detection framework, anomalous data instances can be identified as the model was forced to encode relevant regularities. The working hypothesis revolves around the assumption that *normal* instances can be better reconstructed from the latent space than *anomalous* ones, a difference that can be characterized a posteriori quantifying the reconstruction error.

The structure of AE models follows the following definition: a set of input data, denoted as *X =* (*x*_1_, …, *x*_*n*_) is fed to the model. The latent variables, *Z*, are outputted by an encoder, *F*(*X*), and inputted in the decoder *G*(*Z*), which is trained to reconstruct 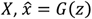. The AE objective is then composed of one term, an unsupervised reconstruction loss [55]:

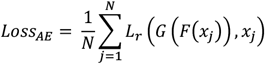

*where N denotes samples, and Lr the reconstruction loss*

#### Normative model development

We trained an AE model with the normative corrected HCP-YA training set (n=1109), composed of 170 tabular features: 68 CT, 50 GMV and 52 WMV features. Afterward, the trained normative model was employed in an anomaly detection framework using the StratiBip external test set. All the code was developed in python using *tensorflow* and *sckikeras* packages.

The first step was to define the best architecture for AE-normative model. The model’s general initial architecture and hyperparameter search space were based on [23]. The model used *selu* activation function and *lecun_normal* weight initialization in all layers [56], except for the last layer of the network that was defined using a *linear* activation function and *gorot* weight initializer. An l2 norm was included in all layers for regularization. The model optimization was based on *Adam* [57] and the loss function, *L*_*r*_, on the *mean squared error*, 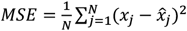. The number of layers, the number of neurons, the batch size, the number of epochs, the learning rate and the l2 norm coefficient were optimized in a 10 fold cross-validation (CV) hyperparameter tunning process with a random search strategy as detailed in the supplementary information. After hyperparameter tunning, the best AE model was retrained on the entire HCP-YA training set.

### 2.5 Normative model application

We applied the trained normative AE model to the external StratiBip test set. From the reconstructed StratiBip data, for each feature, we extracted the RE, the squared error between the original and reconstructed instances, 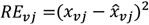. Then, the RE values were integrated with the mean squared error (MSE) for computing the subject’s mean deviation scores (MDS) by averaging the squared error across all the features: 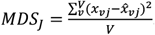. To assess model robustness and variability to training data we employed a bootstrap with replacement strategy. The HCP training set underwent a random selection with replacement for 1000 iterations. Each time, an AE normative model was trained with each bootstrap sample and applied to the StratiBip test set. The MDS values resulting from the 1000 bootstraps were subject to the analyses described in the group-level analysis section – BD-deviating brain features. We computed the percentile 95% confidence intervals in order to evaluate the variability of model performance and extract statistically significant deviating group-level features, in the BD group.

#### 2.5.1 Group-level analysis

The RE metrics (both RE and MDS) extracted from HC and BD individuals of the StratiBip test set were entered in the following group comparisons.

##### 2.5.1.1 BD-deviating brain features

*AE-based anomaly detection*. To assess whether BD individuals differed from HC in terms of their deviation outcomes from the AE normative model, group-level BD vs. HC comparison of feature-RE values was performed.

First, the median MDS between HC and BD subjects were compared. Then, feature-specific RE distributions were compared to identify region-based brain deviating patterns at the group level. For each brain morphological feature, we compared the RE non-normal distributions between HC and BD, using a one-tailed Mann-Whitney U (MWU) test (alternative hypothesis: BD group median to be higher than the HC group), assigning a critical level of 0.05 (uncorrected), and computing the cliff’s delta effect size to quantify the magnitude of the differences. The initial significance criterion was established by evaluating the p-value 95% confidence interval (CI), accepting all tests with a mean p-value bootstrap estimate of less than 0.05. For the features identified from this initial assessment, the effect size was subsequently evaluated and considered significant if its 95% CI excluded zero [24]. The features resulting from this second-level assessment were identified as having significant increase deviations in the BD group.

###### Mass-univariate analysis

We performed a standard mass-univariate analysis to facilitate the interpretation of findings regarding the BD normative deviating brain features results from the previous section. Consistently with our pipeline, the corrected features used in this analysis were the same fed to the AE-normative model. A two-tailed MWU test mass-univariate analysis was employed to assess differences between the distributions of the original corrected feature sets between BD and HC group. The critical level was set to 0.05 and a Bejamini-Hochenberg false discovery rate (FDR) correction was employed for multiple comparisons.

###### MDS-based discrimination of BD vs. HC: ROC curve analysis

Following, we evaluated whether the resulting brain deviations, quantified through the MDS, could discriminate the two StratiBip groups. Each subjects’ REs was summarized with the MDS and an receiver operating characteristic (ROC) curve analysis was employed. The area under the curve (AUC) of the ROC curve was extracted and the optimal discriminative MDS threshold was identified.

##### 5.2.1.2 RE patterns heterogeneity

After assessing group differences we investigated RE patterns heterogeneity within and between groups. We computed the pairwise feature RE absolute differences between every two subjects, in each group separately and then between groups. Then, we summarized the overall results feature-wised with the mean heterogeneity, Eq. 4, where *v* stands for feature, *j1* and *j2* denote two subjects from the same group with N total subjects, and *m* a selected subject from a different group with *M* total subjects. The more the RE outcomes varied across subjects for a specific brain feature, the higher the average difference and the heterogeneity.

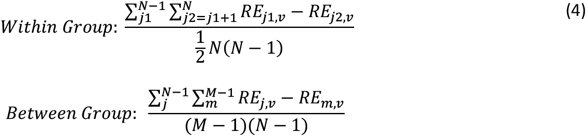

##### 2.5.1.3. RE extreme deviations

Afterwards, we moved away from the description of group central tendencies, i.e., comparing medians/mean, and exploited extreme value statistics concepts to investigate the profiles of the RE distribution tails. First, a leave-one-out (LOO)-CV was performed to extract unbiased reconstructions for all HCP-YA training set subjects. In each fold, all subjects except one were used to train the normative AE model. The left-out subject was used as test sample and its reconstruction was extracted. Then, we applied a block maxima approach, where a series of independent observations are summarized by its maximum value within a specific block [58]. In our case, in each group, each feature was considered a block of data with N independent subjects’ measurements and was summarized with the top 1% mean of extreme values (MEV), i.e., the 99% trimmed mean, Eq. 5, where *k* is the number of data points corresponding to the top 1%. We assessed differences in terms of MEVs for each feature in the three groups, StratiBip HC and BD, and HCP-YA.

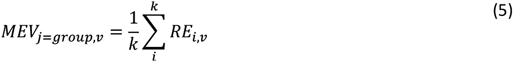

#### 2.5.2 Personalized brain deviating maps

##### 2.5.2.1 Modified z scores

The most promising application of the proposed AE normative modelling framework is to move from group-level to individualized analyses. We charted the StratiBip test set features REs by comparing them with the distributions extracted from the HCP-YA training set with the LOO-CV analysis, via modified z scores, *mZ*. The mZ scores accounts for the median and median absolute deviation (MAD) and is more robust than its parametric version for outlier identification when the underlying data distribution is non-normal [59]. Besides, MAD is a robust measure that captures the dispersion around the median while not being influenced by extreme values and the range of the dataset. First, analysing the HCP-YA normative RE outcomes, we calculated the RE median for each feature, *E[RE*_*HCP;v*_], which we considered as the expected model normative RE. Then, we calculated the MAD, the measure of model uncertainty for reconstructing feature *v*, adjusting the MAD with a correction factor of 1/Q(0.75), where Q(0.75) corresponds to the 75^th^ quantile in the respective normative feature distribution [59]. Then, the mZ score foresees that each new data point be standardized with the median and MAD of the normative expected RE distribution, Eq. 6, and was used to compute personalized deviating brain maps for each subject in the StratiBip test set. Afterwards, we defined an abnormality criterion based on the MAD, to derive abnormal features at individual level. Usually, when data is normally distributed, a known threshold for outlier detection is the measure of 3 standard deviations, or 3.5 MADs [60], [61]. In our case, we defined a threshold for each feature based on its specific normative RE distribution. Our data did not follow a normal distribution and we assume that each feature was encoded differently by the model, having different expected normative RE outcomes. Thus, we translated this feature-specific encoding into a definition of feature-specific abnormal thresholds. For each normative RE feature distribution, we took the mZ threshold corresponding to the 99^th^ percentile. Thus, an individual feature was considered abnormal if fell in the top 1% of the normative RE expected distribution.

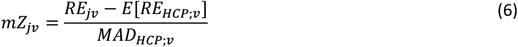

##### 2.5.2.2 Spatial overlapping deviating patterns

Finally, we investigated the spatial overlap of deviating brain maps within groups. First, for each feature, we computed the frequency of abnormality occurrences within each group. Next, the subjects’ brain deviating maps were transformed into descriptive sets of abnormal features, and the pairwise subject overlap coefficient (OC) and Jaccard similarity (J) were computed within and between groups, Eq. 8, 9. The OC calculates the minimal overlap between two item sets, ranging between 0 and 1, where 1 is totally similar or one set is a subset of the other, Eq. 7. On the other hand, the Jaccard coefficient calculates the total similarity between two item sets, ranging from 0 to 1, where 1 stands for totally similar., thus testing whether two sets share the same members, accounting for all the members, Eq. 7.

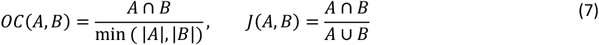

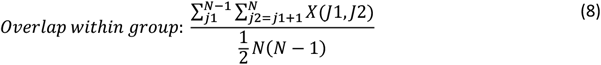

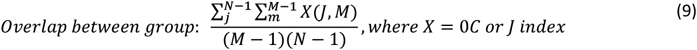

## 3. Results

### 3.1 Datasets Characteristics

The normative training set was extracted from the HCP-YA sMRI dataset [28], with 1109 subjects (median age = 29.00 years, 604 females, 505 males), whereas the external test set was derived from 550 subjects, 363 HC (median age= 27.00 years, 189 females, 174 males) and 187 subjects with BD (median age= 30.00 years, 101 females, 86 males), from 7 acquisition sites of the StratiBip network (Table S1 and Fig. S1). All included subjects were between 22 and 37 years old. A Kruskal-Wallis test revealed significant age differences among the three groups, HCP-YA, StratiBip-HC, and StratiBip-BD (χ^2^(2)= 34.85; p<10^−7^); the post-hoc comparisons showed that StratiBip-HC were younger than HCP-YA and StratiBip-BD subjects (Table S1). On the other hand, based on a Chi-Square test of independence, no significant differences among the three groups were found for sex proportions (χ2(2, N=1659)=0.6338, p=0.728). More detailed information on the sample characteristics in each site can be found in Table S2.

### 3.2 Multi-site harmonization effectiveness

We checked the quality of site effect removal performed via M-ComBat application. Before harmonization, all feature set distributions (GMV, WMV, CT) for the HCs among the 8 sites (HCP site and 7 StratiBip sites) resulted significantly different (p<1e-29) but no differences were detected among sites after harmonization (p>0.680).

For BD in the 7 StratiBip sites, all feature sets were significantly different across sites (p<1e-12) before harmonization, whereas statistically significant differences remained for CT and GMV features (p<0.018) after harmonization; the pairwise post-hoc comparisons corrected for multiple tests showed that differences survived for CT features between site 4 and site 6 (Table S3 and Fig. S2). A second quantitative check was performed by probing how the harmonization affected a support vector machine (SVM) model trained to classify sites based on the entire feature set. A substantial decline in average F1 score was observed, from 95% before harmonization to 23% after harmonization, and all sites showed a decrease in F1 score to below chance-level (Table 1). Group- and feature set-specific SVM site classification results were also extracted (Table S4). Further analyses assessing M-ComBat performances in terms of biological effect preservation were performed (Fig. S3, Tables S5-8).

**Table 1.** F1-score SVM site classification before and after harmonization.

| F1_score | HCP-YA<br>(N=555) | 1<br>(N=38) | 2<br>(N=82) | 3<br>(N=51) | 4<br>(N=14) | 5<br>(N=41) | 6<br>(N=33) | 7<br>(N=22) | Weighted<br>Average<br>(N=836) |
| --- | --- | --- | --- | --- | --- | --- | --- | --- | --- |
| Before<br>Harmonization | 1.00 | 0.88 | 0.89 | 0.94 | 0.38 | 0.99 | 0.72 | 0.67 | 0.95 |
| After<br>Harmonization | 0.30 | 0.03 | 0.08 | 0.12 | 0.00 | 0.15 | 0.12 | 0.09 | 0.23 |

### 3.4 AE-based normative model performance

When trained on the HCP-YA training set, the AE normative model achieved a training loss MSE of 0.182 ([0.179;0.185]; 95% CI) and a validation loss MSE of 0.222 ([0.211;0.233]; 95% CI) after 2000 training epochs (Fig. S4). After training, we extracted the AE model reconstructions for the StratiBip external test set data and computed the respective REs and MDS by subject, by group, and by feature-by-group. Concerning the subjects’ MDS, as expected, the BD group showed a significantly higher MDS median, 0.2264 ([0.2210,0.2324]; 95% CI) compared to the HC group, 0.1988 ([0.1945,0.2030]; 95% CI). Such difference was statistically significant since the CI for the two groups did not overlap, or, in other terms, the median MDS difference CI did not include zero, -0.02760 ([-0.03390, -0.02155]; 95% CI). The feature-wise MDS 95% CIs are reported in Fig.S5.

### 3.5 Group-level BD vs. HC comparisons

#### 3.5.1 BD-deviating brain features

We employed the trained AE model to extract the StratiBip external test set reconstruction errors and calculated the respective MDS. Several features from all types (CT, GMV, WMV) were found to have significantly higher deviations in the BD group, identified by a significant Cliff’s delta effect size and an uncorrected bootstrap mean estimate pvalue <0.05 (Fig. 2). We identified higher BD deviations in CT in the right inferior temporal gyrus, and in volumes of subcortical and adjacent regions belonging to the cerebellum and the limbic system (hippocampus, striatum, globus pallidus). To provide a reference for the AE model findings, we also performed a standard mass-univariate statistical BD vs. HC comparison using a two-tailed MWU test (p<0.05; uncorrected and FDR corrected). Only the WMV surrounding the left globus pallidus emerged as significantly different after correcting for multiple tests (Table S11).

**Fig 2.**
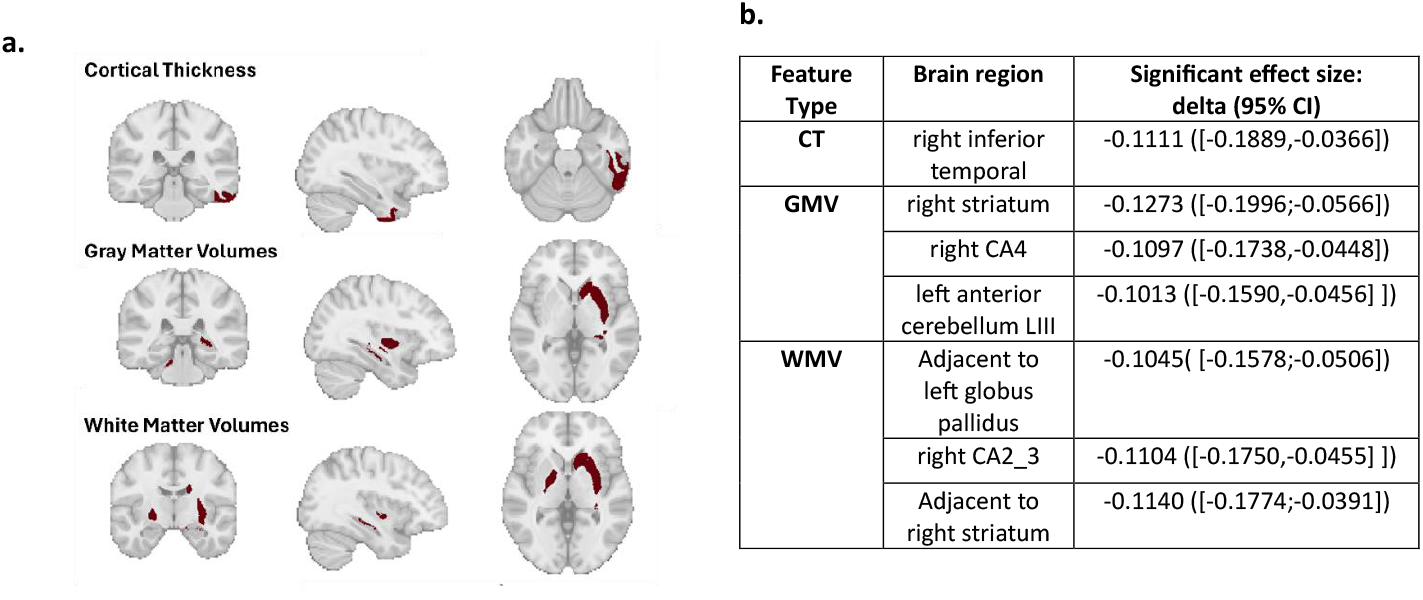
Brain features with significantly higher deviations in BD. a, Representation of all brain features that were identified with significantly higher deviations in the BD group. **b**, Table describing the identified features and their associated 95% CI cliff’s delta effect size.

#### 3.5.2 RE patterns heterogeneity

We then quantified the feature-wise RE heterogeneity within and between each group by computing the average RE differences across pairs of subjects (Fig. 3). In general, RE patterns were more homogeneous in the HC group, with a maximum mean pairwise difference of 0.59 ± 1.2, compared to 1.8 ±6.8 in the BD group. Overall, for both groups, CT and WMV features presented higher levels of heterogeneity than GMV features. Among all features, the WMV of the left and right Stratum displayed the highest pairwise RE difference among BD subjects, ranking 1^st^ and 2^nd^ in terms of heterogeneity (Fig.3a), but not among HC subjects, ranking 6^th^ and 11^th^ (Fig.3b); of note, these features showed the highest group difference, i.e., the absolute pairwise difference between subjects’ RE from the two groups, ranking 1^st^ and 2^nd^ (Fig.3c). In the BD group, other features with high RE heterogeneity included WMV of the left alveus, left HCA1, left and right CA2_3 and left CA4, and CT of left para-hippocampal gyrus and bilateral medial orbitofrontal cortex. In the HC group, the WMV of left CA4 and anterior cerebellum displayed the highest heterogeneity, followed by the left alveus, right CA2_3, left CA2_3, and left Stratum. Apart from WMV in the left and right Stratum, the features differing the most in terms of RE magnitudes between HC and BD groups included WMV in left alveus and CA4, bilateral CA2_3 and CT of para-hippocampal gyrus.

**Fig 3.**
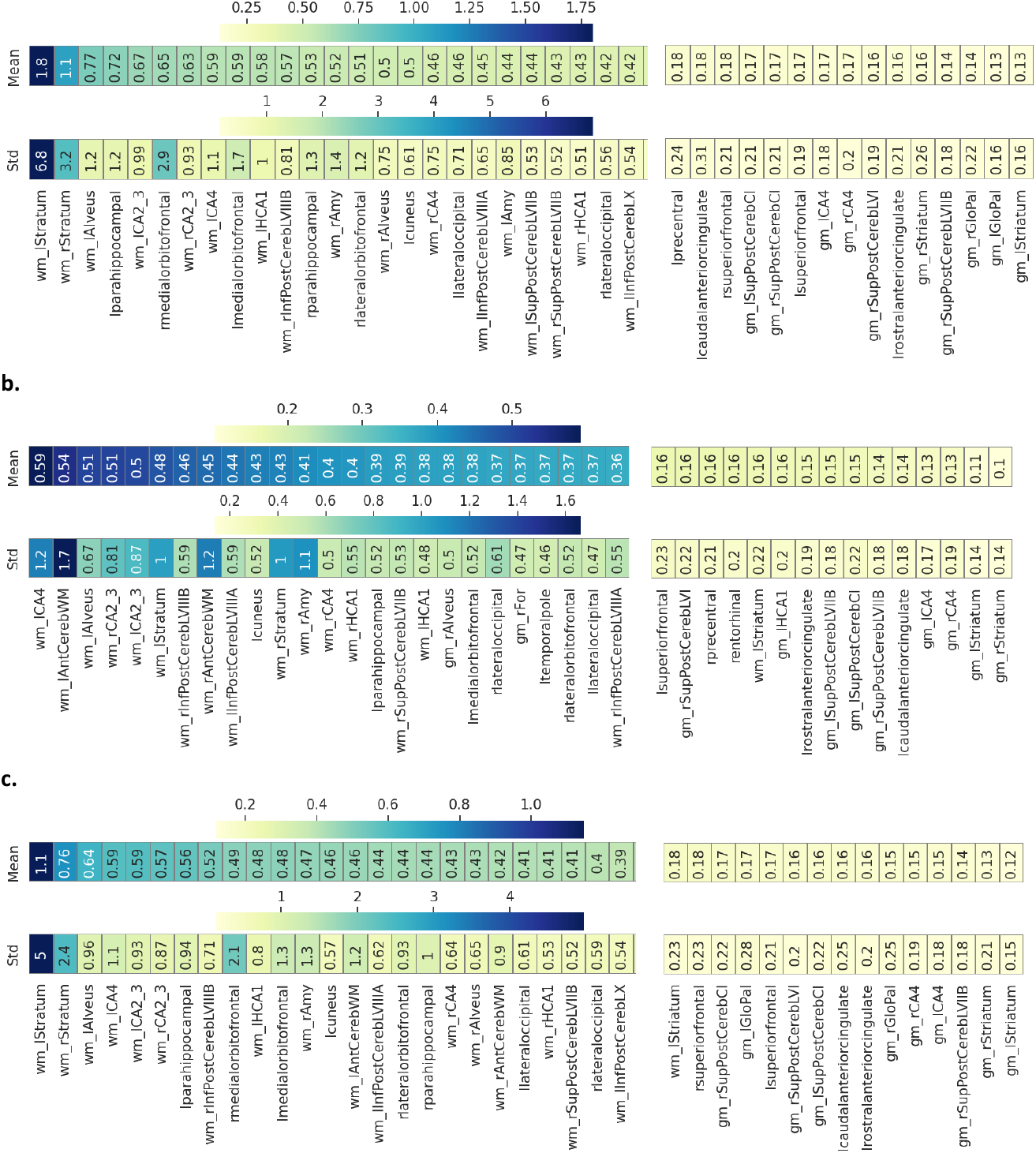
RE heterogeneity within and between groups. The mean and standard deviation RE pairwise differences are shown in a sorted heatmap, including 25 features with the highest heterogeneity levels and the least 15, for **a**, BD group; **b**, HC group; **c**, between the two groups HC vs.BD.

#### 3.5.3 RE extreme deviations

We then modeled extreme REs applying a block maxima approach, where each feature was summarized by its extreme values within each group (HC, BD, HCP-YA). Employing a LOO-CV strategy, we retrieved unbiased reconstructions for each subject in the normative HCP-YA training set and constructed a normative RE distribution for each feature.

Including only the top 1% REs (99% trimmed), we compared the MEV between the normative HCP-YA training set and StratiBip HC and BD test sets (Fig.4). In WMV and CT feature sets, the overall maximum MEV in the normative group resulted lower when compared with the 2 StratiBip groups; conversely, all GMV features in the StratiBip HC group resulted within the respective normative group range. In all feature sets, selected features showed MEV differences among the three groups. In general, the BD group was characterized by a more pronounced extreme value profile, resulting in 7 CT, 4 GMV, and 4 WMV features with at least a double MEV compared to the normative and the StratiBip HC groups (Table 2). In contrast, in the HC group, only 2 WMV features showed at least a double MEV compared to both the normative range and BD group (Table 3).

**Table 2.** Summary of features with at least a double MEV in the StratiBip BD group vs. others.

| Features MEVs |  | BD-StratiBip | HC-StratiBip | HCP-YA |
| --- | --- | --- | --- | --- |
| CT | Left parahippocampal gyrus | 6.9 | 3 | 3 |
|  | Right parahippocampal gyrus | 8.7 | 2.7 | 2.1 |
|  | Right lateral orbitofrontal | 6.9 | 3 | 2.8 |
|  | Left medial orbitofrontal | 11 | 2.8 | 2.4 |
|  | Right medial orbitofrontal | 15 | 1.9 | 2.7 |
|  | Right superior parietal | 6.3 | 1.4 | 1.7 |
|  | Right rostral anterior cingulate | 4.9 | 1.8 | 1.5 |
| GMV | Right Inferior posterior CerebLVIIIA | 4.2 | 1.9 | 1.2 |
|  | Adjacent to left Fimbria | 2.4 | 1.2 | 1.2 |
|  | Left Thalamus | 6.3 | 2.5 | 1.2 |
|  | Right Thalamus | 5.3 | 2.5 | 0.75 |
| WMV | Right Stratum | 19 | 6 | 5.3 |
|  | Left Stratum | 44 | 5.9 | 3.6 |
|  | Left HCA1 | 5.6 | 2.6 | 2 |
|  | Adjacent to right Thalamus | 4.7 | 2.3 | 1 |

**Table 3.** Summary of features with at leat a double MEV in the StratiBip HC group vs. others.

| HC group features |  | BD-StratiBip | HC-StratiBip | HCP-YA |
| --- | --- | --- | --- | --- |
| WMV | Left Cerebellum | 2.5 | 10 | 1.4 |
|  | Right Cerebellum | 2.9 | 5.8 | 1.3 |

#### 3.5.4 BD vs. HC MDS-based discrimination

We assessed whether the subjects’ MDS would enable the discrimination between the BD group and HC one in the StratiBip test set, achieving an AUC-ROC of 0.6129 ([0.5989, 0.6270]; 95% CI). The optimal MDS threshold to differentiate HC vs. BD was 0.2138 ([0.2096,0.2181]; 95% CI) which yielded a mean accuracy of 58.3% ([56.4%;60.4%]; 95% CI). Then, we inspected whether accounting for extreme value statistics would enhance this discrimination. This time, each subject was summarized by its extreme values under a block maxima approach, with the MEV (99% trimmed). Then, the ROC curve analysis was repeated, obtaining an AUC-ROC of 0.6218 ([0.5999, 0.6452]; 95% CI), for an optimal MDS threshold to differentiate HC vs. BD of 1.9032 ([1.8417,1.9723]; 95% CI) yielding a mean accuracy of 59.0 % ([56.2%;61.8%]; 95% CI), a slight improvement when compared to using central tendency statistics to summarize the RE outcomes, i.e., the MDS.

### 3.6 Personalized brain deviating maps

Individual brain deviations were also employed for subject-level statistical inference. We calculated the mZ for the StratiBip dataset using the HCP-YA feature-wise median and MAD. Then, for each feature, we retrieved the 99^th^ percentile in the normative HCP-YA distribution and used it as the normative mZ threshold, enabling the identification of subject-level deviating features (mZ > 99^th^ percentile) for each StratiBip individual (Fig S6). We report the resulting brain CT, GMW, and WMV deviating maps of two exemplar subjects from the StratiBip test set, one control and one affected with BD (Fig. 5). The mZ distributions of all features in the StratiBip HC and BD groups are reported in Fig. S7.

**Fig 4.**
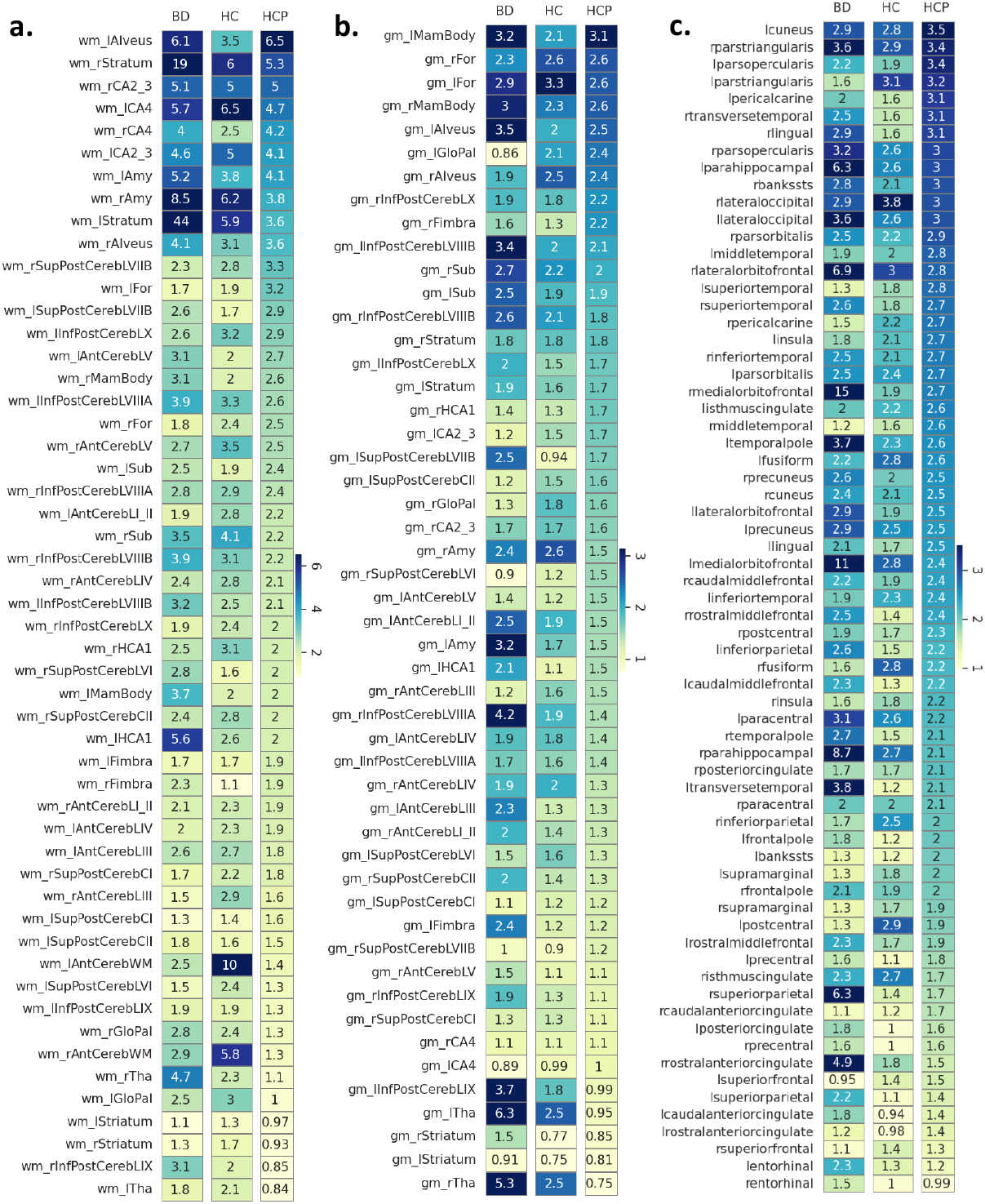
Feature MEVs in the normative HCP-YA and StratiBip BD and HC test set groups. In the top heatmaps (**a**. WMV features, **b**. GMV features, **c**. CT features), the feature-wise MEVs for the StratiBip BD (BD column), StratiBip HC (HC column) and normative HCP-YA (HCP column) groups are plotted. Features are sorted in descending order based on the normative HCP-YA MEVs. The StratiBip BD and HC group heatmaps are color-coded in the same range as the normative HCP-YA one to highlight deviations from the normative expectation within the same brain feature.

**Fig 5.**
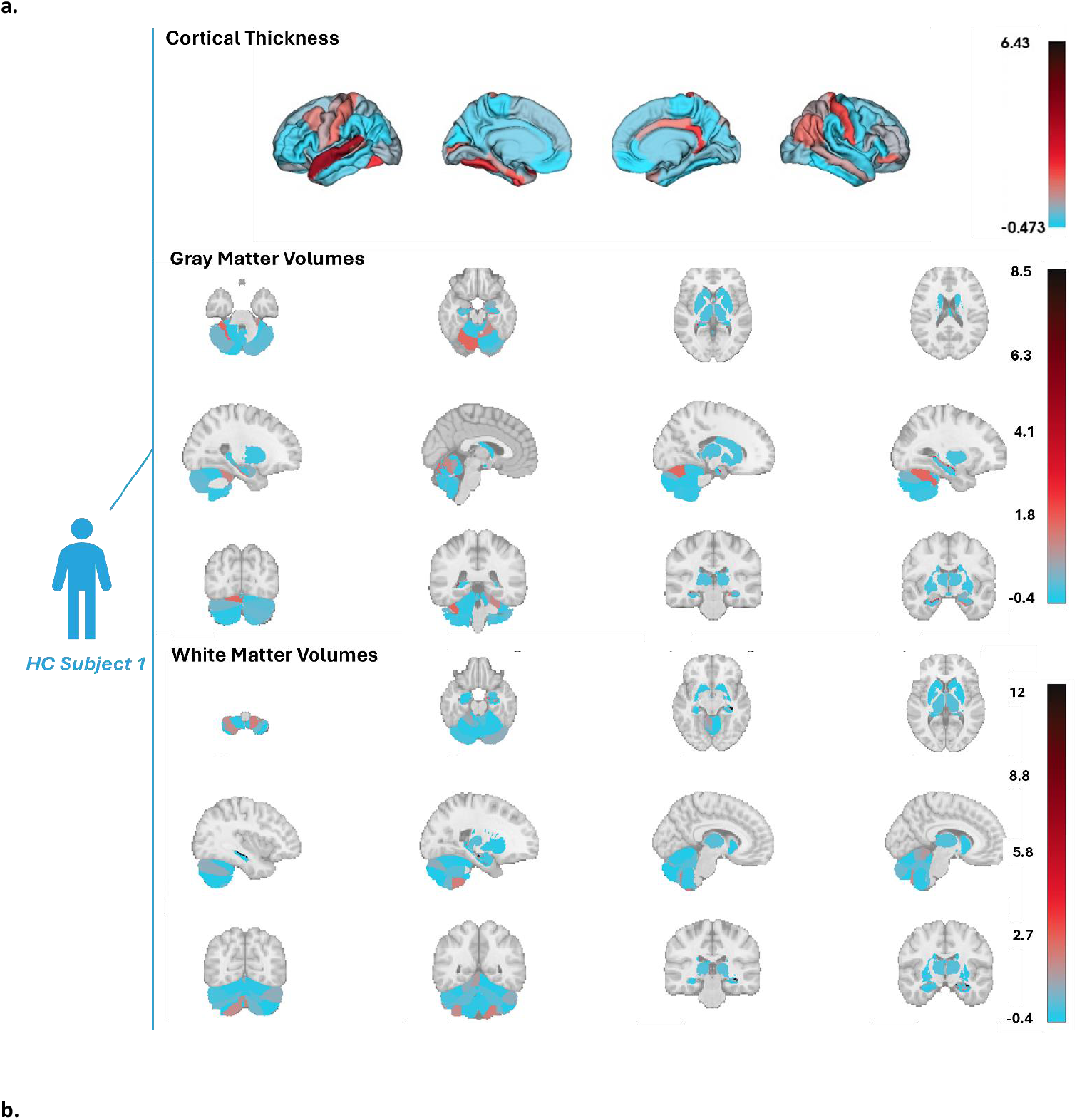
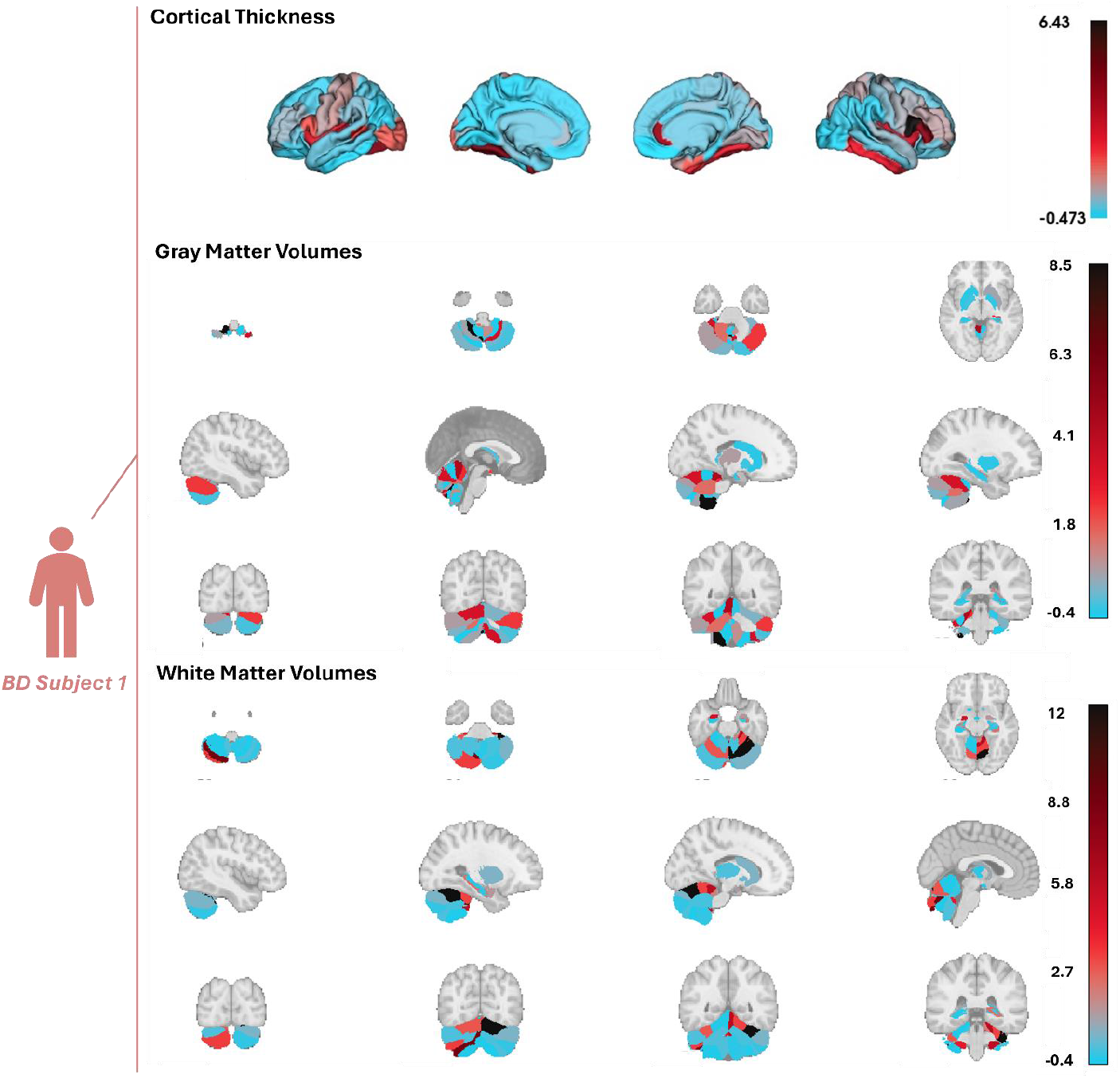
Individual deviating brain maps. We plot the deviating CT, GMV, and WMV feature maps for 2 subjects: **a**, HC subject CT, GMV and WMV mZ scores; **b**, BD subject CT, GMV and WMV mZ score. The color bar range is shared between the two subjects for each feature set group to better highlight differences in the deviating maps.

Next, for the two StratiBip groups, we inspected the prevalence of subject-level abnormal features. Across all feature sets, subjects belonging to the HC group had an average of 1.3% abnormal features, corresponding to about 2 features per subject, while in the BD group, this average percentage increased to 1.9%, corresponding to 3 abnormal features per subject. For each feature, we inspected the percentage of abnormal occurrences for each group (Fig. 6). In the BD group, the highest prevalence of abnormal patterns (11% of subjects) was found for the WMV adjacent to the left globus pallidus, followed by the GMV of the right thalamus (7.5%) and WMV: of right inferior posterior CerebLIX, surrounding the bilateral thalamus, of left HCA1 and right inferior posterior CerebLVIIIB (7%). Of note, in the HC group, the highest frequency of abnormal cases was also observed for the WMV adjacent to the globus pallidus (6.9%), followed by GMV of right thalamus (6.3%), WMV of left anterior Cerebellum (6.1%) and adjacent to bilateral thalamus (5.5%), and GMV of right amygdala (5.2%). The intra-group and inter-group similarity was also assessed by employing the average pairwise overlap coefficient (OC) and the Jaccard similarity index (J), achieving (i) in the HC group, higher level of similarity compared to the BD group, (ii) in the BD group, lower level of similarity compared to the inter-group one (BD-HC) (OC_HC_=0.72; OC_BD_=0.60; OC_HC_BD_=0.67 | JC_HC_=0.32; JC_BD_=0.23; JC_HCvs.BD_=0.27).

**Fig 6.**
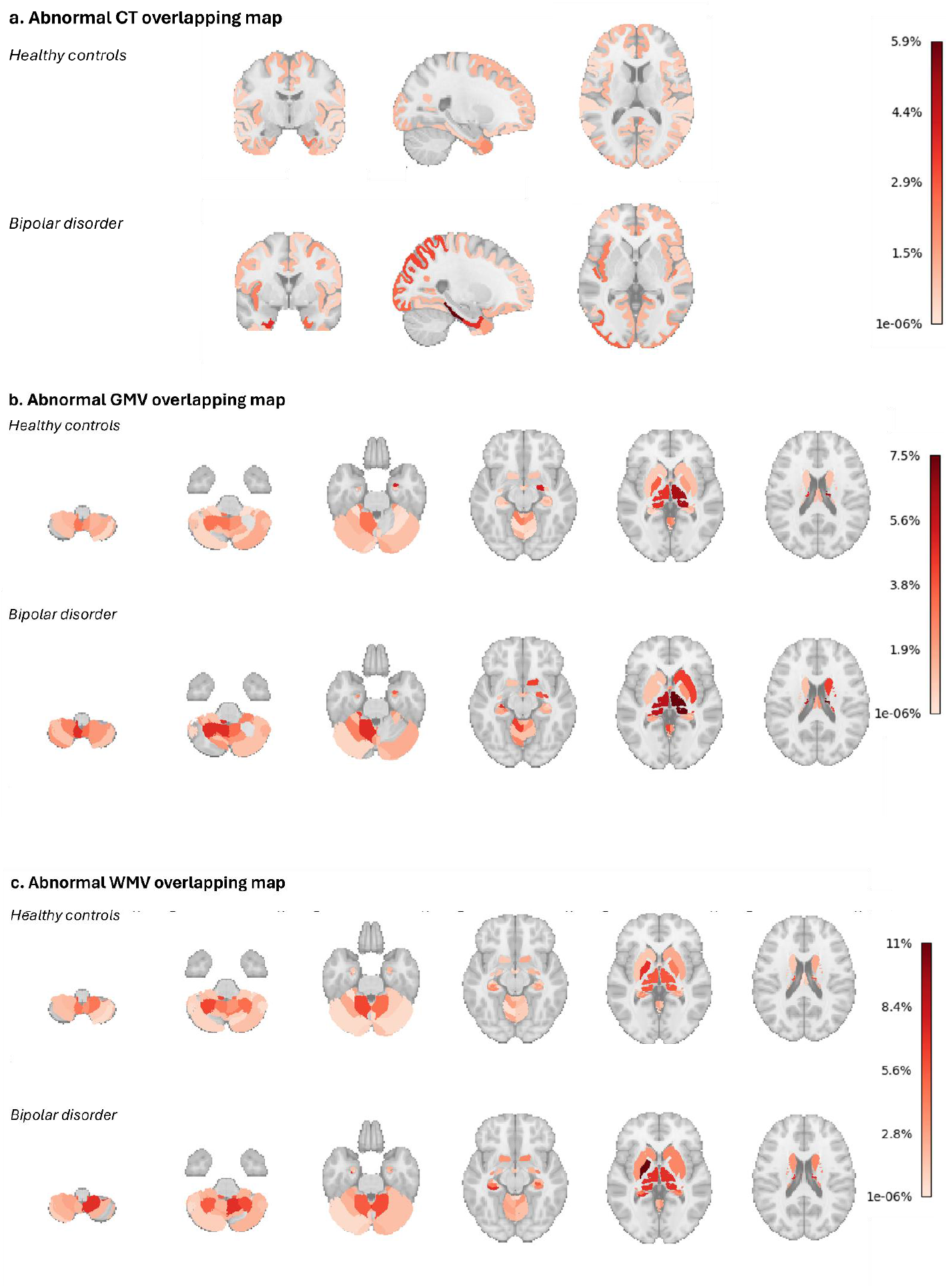
Feature set percentage of abnormalities for each group. For each feature set and brain region, the brain maps show the prevalence of individuals, in percentage, with abnormal features within each group, HC and BD.

## 4. Discussion

In this study, we designed a robust, generalizable, and extendable end-to-end pipeline for brain morphological multivariate normative modelling and personalized anomaly detection based on deep AEs. The pipeline embodies data pre-processing fully integrating training with external validation (including data harmonization and biocovariates confounders removal), normative modelling, and statistical comparison steps. This innovative framework was used for performing personalized and group-level statistical inference on brain morphological deviations characterizing individuals with BD. The AE-based normative model was built on brain regional features from the large normative HCP-YA cohort and tested on features from the external multi-site StratiBip cohort, including both controls and BD individuals.

First, we showed the effectiveness of the proposed pipeline in removing site-related effects from the external StratiBip test set. This allowed us to leverage and integrate external test sets acquired in different sites, enabling robust comparisons and increasing statistical power. Then, we proved the effectiveness of our approach in characterizing brain morphological deviations in test samples, identifying subject- and group-level tendencies but also heterogeneity and extreme deviation patterns within and between groups.

Our findings showed that, on average, group-level deviations from the norm were higher in BD compared to HC; in the BD group, RE patterns were also more heterogeneous and with greater extreme values than in HC. Moreover, at the individual level, the most prevalent deviations were observed in features that were common between HC and BD, but such prevalence was increased in the BD group. Notably, we also found that the spatial overlap of individual-level brain deviating maps was greater between BD and HC subjects than within the BD group itself.

The latter evidence is in line with the hypothesis that brain morphological alterations in BD, and in general in psychiatric disorders, are subtle and might be nested within the spectrum of normative interindividual variability. Overall, these results support the conceptualization of BD as a non-unitary disease with a variety of neurobiological dimensions, whose characterization paves the way to the identification of personalized signatures of disease and more effective interventions.

### Innovative features of deep normative modelling and anomaly detection framework

In this ever-growing research context, the proposed pipeline is distinguished from previous ones by combining the following features. Although AEs have been proposed before in literature for brain normative modelling [23], our differs due to the innovative inclusion of a generalizable confounder removal step in the DL normative modelling pipeline, which enabled its effective translation to external datasets. In S. Rutherford *et al*. [62] authors proposed a method to extend a pretrained Bayesian regression model to data from new sites, but site-related variation is modelled together with features-of-interest within a single regression model by including the site variable as covariate, impending its usage in our deep learning framework. Just as innovative is the multivariate nature of deep learning AEs for normative modelling. AEs are suitable for integrating data and have been successfully applied to multimodal datasets [63]–[65]. In the search for brain markers of BD, our study is the first to employ a multivariate normative analysis framework that integrated CT, GMV and WMV features for the subject- and group-level characterization of this complex disease.

### CR pipeline was effectively applied to external datasets

To our knowledge, our study is the first to embed in the normative and external validation framework the removal of site-related and biological confounding effects from the brain features, in a deep learning normative model application. We considered working with confounder-free data as a pre-requisite towards more interpretable DL models. The inability to know what information drives the performance of a ML model can lead to erroneous result interpretations, known as source ambiguity problem [26], [53], [66]. Therefore, to achieve more interpretable models, it is recommended to control for alternative sources of information from the target of interest, known as covariate adjustment or confounding-effects correction. In our study, we showed that the normative M-ComBat successfully harmonized the external StratiBip test sets with the HCP-YA training set. This ComBat variation has been shown to effectively harmonize data from different sites and has been recently employed in a multi-site PET study in an external validation framework [50]. Moreover, our confounder removal pipeline was developed in a normative framework; therefore, in both harmonization and biocovariates linear regression models, any associations between diagnosis and brain features were not modelled, as this could have led to data leakage problems and consequently biased the model [53], [54]. The normative site and biological confounding effects were assumed to be generalizable to patient data, and both estimated effects (ComBat parameters, biological beta coefficients) were applied equally to data from the StratiBip HC and BD groups.

### AE-based normative modelling empowered identification of group-level brain morphology deviations in BD

Group-level analyses on brain morphological correlates of psychiatric disorders have been extensively performed in literature, but only few in terms of normative deviation metrics [14], [20], [22], [23], [67]. Since normative models can detect individual deviations from the norm, they are especially suitable for unravelling brain heterogeneity in BD. Our findings showed higher median deviations in BD compared to HC; specifically, volumes of the basal ganglia and adjacent to it (striatum and globus pallidus) and from the hippocampus (CA4, CA2_3) revealed increased deviations in BD compared to HC. The WMV surrounding the globus pallidus was also significant in the mass-univariate case-control analysis that we used as reference, supporting the neurobiological plausibility of the AE-based normative findings. These group-level deviations are in line with existing literature on BD, suggesting morphological alterations in brain regions involved in affective processing, including the basal ganglia, hippocampus, and temporal regions observed in our study. In the case-control mega-analyses of the ENIGMA BD Working Group, BD was found to be associated with cortical thinning in inferior temporal regions and with volumetric reduction in the hippocampus [4], [10]. Additionally, in another study employing a univariate normative approach, individuals with BD were also reported to have GMV deviations in cerebellar and temporal regions [14].

While the overall agreement with the existing evidence supports the reliability of the deviations observed in our BD sample, it should be considered that our multivariate findings reflect hidden links among the brain features. Due to the multivariate nature of deep AEs, the features that emerged as deviant should be understood as patterns of alterations rather than region-specific alterations.

Regarding the BD group discrimination, the whole-brain MDS presented a low discriminative power when compared with the state-of-art, achieving an AUC-ROC of 0.61 and an accuracy of 58.3% using the best MDS threshold. A recent review on machine learning studies that attempted to classify BD vs. HC reported a range of prediction accuracies of 59%-78% based on WMV and GMV predictors [68] in parallel, the ENIGMA BD Working Group reported an AUC-ROC of 0.7149 (0.6939-0.7359) using cortical thickness, surface area and subcortical volumes; this improved performance could be due to different factors, like the inclusion of a bigger BD sample or the non-removal of biological effects from the brain features used for classification [69].

### Distribution and extreme pattern analyses highlighted brain morphology heterogeneity in BD

Our normative model was exploited to assess and compare the heterogeneity and extreme profiles of the feature deviating patterns in BD and HC groups.

BD individuals presented higher levels of heterogeneity, especially for WMV in subfields of the hippocampus, alveus, and cerebellum, and for CT of parahippocampal and medial orbitofrontal regions. The highest difference between groups, highlighting much greater heterogeneity in BD, was found for the WMV of the bilateral stratum. This more marked heterogeneity of REs reflects a greater model variability in reconstructing the data, which in turn is suggestive of brain morphological heterogeneity in the BD group. The enhanced brain heterogeneity could underlie the phenotypic variability of individuals affected by BD, which represents one of the main reasons that so far have impeded the identification of objective brain markers of disease [70]. In this respect, an increasing body of evidence is remarking the need to adopt a dimensional perspective for identifying the brain endophenotypes of clinical dimensions that are shared between BD and other disorders in the psychotic or affective spectrum [71], [72].

Interesting evidence on BD was provided by the assessment of extreme deviations; Our findings suggest more pronounced extreme deviations in BD, being characterized by the greatest number of features with a MEV that was more than the double of both StratiBip HC and HCP-YA groups. Moreover, the discrimination between the BD and HC groups improved when using MEVs instead of MDS as subject-level deviating scores, achieving an AUC-ROC of 0.62. This suggests that examining extreme values can enhance the separability between groups.

High extreme deviations were found in features that showed marked heterogeneity in the BD group, including WMV of bilateral Stratum and left HCA1 and CT of left parahippocampal and bilateral medial orbitofrontal regions. We hypothesize that this heterogeneity was driven by the incidence of extreme values in these features, possibly reflecting pronounced phenotypic differences in the BD group. Notably, in [30] authors identified normative deviation scores of the GMV on the left middle orbital frontal gyrus as the most reproducible feature to discriminate BD from HC, applying a random forest classifier. We hypothesize that only a sub-group of subjects present more severe alterations in these regions and this may drive both the increased heterogeneity/extreme values observed in the present results and the high discriminatory stability in the second.

### AE-based normative modelling empowered the creation of personalized brain deviating maps

Our normative framework allowed us to characterize subjects at the individual level. Individual brain deviation maps were built employing the mZ scores and considering a conservative 99^th^ percentile threshold for abnormality. On average, subjects affected with BD and HCs showed a similar percentage of abnormalities, slightly higher in BD (1.9%) than in HC (1.3%). The maximum spatial overlap of features identified as abnormal was identified for the WMV surrounding the globus pallidus, expressed in 11% of BD subjects and in 6.9% of HCs, followed by the GMV of right thalamus (7.5% in BD, 6.3% in HC). Interestingly, previous univariate normative studies on BD reported the highest spatial overlap of abnormalities in the thalamic region, showing around 2% in [14], and 5.17%-8.19% in [73], and high discriminatory stability of GMV thalamus deviations [30]. Overall, our results show that abnormalities in BD spread mostly through the volumes of the bilateral thalamus and adjacent to it, hippocampus subregions, and cerebellum.

Of note, this personalized inference on BD subjects unravelled brain morphological abnormalities in regions that did not emerge from the group-level comparisons. These regions included the thalamus, for which volumetric alterations have been previously reported in case-control mass-univariate comparisons [4]. It should be noticed that thalamic volume was deviating in a number of HC and BD subjects, albeit with higher frequency in the last group. This might be attributed to thalamic alterations being nested in healthy variations, overcoming this expected variability only for a subset of patients.

Overall, across all features, we found a lower overlap of individual abnormalities in BD than in HC. In the BD group, a minimal subset of abnormal features replicated on average at the pairwise abnormal spatial maps comparisons (OC=60%), but the complete overlap of deviating patterns was lower (J=23%). Noticeably, abnormal profiles of BD subjects overlapped more with other HCs than with other BD subjects. These results further asserted the heterogeneity of BD and are in agreement with the accumulating evidence that brain changes in BD, as in other psychiatric disorders, might be nested within healthy variations [20], [74].

## 5. Limitations

Several limitations of this study should be highlighted. From a clinical perspective, only young adults were included, which prevented us from performing a more comprehensive analysis of BD in the entire lifespan. Additionally, in the StratiBip test set, sample size and biological covariates were not equally distributed among sites, and this could have affected the results. Dataset diversity and numerosity should be increased in future works to create a more generalizable normative framework inclusive of all age ranges. Another limitation concerned the adjustment for biological covariates. Data was not corrected for medication on BD, as this was not considered in the biological covariates modelling, therefore we cannot exclude that the significant group differences and brain deviations might be driven by medication effects. Similarly, we did not account for comorbidities which might be important to distinguish between disorder-specific effects and others.

Other limitations concern the implemented methodology. Although using confounder-free data contributes to the development of more interpretable DL models, Combat and biological covariates linear regression have their own caveats in terms of confounding source modelling. The former relies on a Bayesian framework for statistical inference of site effects and estimates might be affected by sample numerosity and imbalance between sites. Second, linear regression, although simple and easy to implement, might not capture completely the biological effects if these encompass non-linearities. Nonetheless, up to this date, there is not a gold standard to deal with confounding effects in neuroimaging to achieve confounder-free data, and both methods are widely employed in literature.

On another note, the uncertainty of estimation of the MRI-based features was not evaluated. CAT12 brain tissue segmentation is based on algorithms that might struggle when encountering small brain regions with mixed tissues (gray and white matter) and borders. For example, subcortical gray matter regions on basal ganglia and thalamus have lower GM-WM contrast, with the high content of cellular iron rendering the T1-w signal similar to that of WM [75]. Due to the higher probability of incorrect tissue segmentation in these regions, we incorporated all volumetric estimates produced by CAT12 based on the CoBra atlas. This approach included both WMV estimates for GM regions, and vice-versa, and were interpreted as the volume adjacent to the respective region. These CAT12 estimates might stem from intrinsic limitations in the segmentation software’s voxel-based tissue classification or poor subject-atlas alignment. By utilizing all volume estimates, we avoid excluding potentially relevant information due to cherry-picking selection. Nevertheless, we cannot exclude that our results might reflect the uncertainty associated with these volumetric estimations.

Lastly, with respect to the normative model, a central limitation of our AE-based normative model concerns the lack of directionality information on deviations. Also, we found abnormal features for both HC and BD groups, showing that encoding normative levels is not straightforward and true normative ranges might encompass and generalize to non-normative data as well. On the one hand, the greater brain deviations in BD did not yield sufficient discriminative performance from a clinical application perspective. On the other hand, these findings remark the need to adopt a dimensional perspective for the personalized assessment of brain and phenotypic characteristics in BD as in the general population. A natural extension of this work would therefore be to perform a complete clinical characterization of the deviating scores and more deeply explore the patients’ stratification.

## 6. Conclusion

In this study, we developed a generalizable end-to-end multivariate normative modelling and anomaly detection framework based on deep AEs. The novelty of our pipeline resides in the integration of data harmonization, biological confounder removal, and integration of CT, GMV, and WMV in a multivariate AE-based normative model in an external validation framework. We demonstrated the successful application of this framework in the search for brain morphological deviations in BD, employing anomaly detection in an external multi-site test set composed of HC and BD subjects on a normative model trained with the HCP-YA cohort. Our findings support the hypothesis that brain morphological alterations in BD are heterogeneous and partly nested within healthy interindividual variations, remarking the importance of moving from categorical diagnoses to a transdiagnostic dimensional perspective. In this perspective shift, our multivariate normative modelling framework could capture individual brain differences that might be used for making more effective and personalized clinical decisions.

## Data availability

The HCP-YA normative dataset is publicly available on connectomeDB platform *https://db.humanconnectome.org)*. The StratiBip dataset is governed by data-use agreements or sponsor restrictions and therefore not publicly available.

## Code availability

The custom code used in this study is available for research purposes (GitHub repository https://github.com/inesws/Normative_AE.git). A demo test code is available that allows to try the trained model with some pre-processed and corrected HCP-YA exemplar data. In order to apply the trained model to new data, researchers should follow the instructions.

## Funding

I.W.S. was supported by grants from ERAINS-Italy, project funded under the National Recovery and Resilience Plan (NRRP), Mission 4, “Education and Research” -Component 2, “From research to Business” Investiment 3.1 -Call for tender No. 3264 of Dec 28, 2021 of Italian Ministry of University and Research (MUR) funded by the European Union – NextGenerationEU, with award number: Project code IR0000011, Concession Decree No. 117 of June 21,2022 adopted by the Italian Ministry of University and Research, CUP B51E22000150006, Project title “EBRAINS-Italy (European Brain ReseArch INfrastructureS-Italy).

E.M. was partially supported by the Italian Ministry of University and Research (grant numbers 2022RXM3H7 and P20229MFRC) and by the Italian Ministry of Health (grant n. GR-2018-12367789).

F.P. was supported by Italian Ministry of Health, Ricerca corrente 2024.

P.B. was partially supported by grants from the Italian Ministry of University and Research (Dipartimenti di Eccellenza Program 2023–2027 -Dept of Pathophysiology and Transplantation, University of Milan), and the Italian Ministry of Health (Hub Life Science-Diagnostica Avanzata, HLS-DA, PNC-E3-2022-23683266– CUP: C43C22001630001 / MI-0117; Ricerca Corrente 2024).

## CRediT authorship contribution statement

I.W.S. Conceptualization, Methodology, Software, Validation, Formal analysis, Writing -Original Draft, Visualization.

E.T. Data Curation, Formal analysis, Writing -Review & Editing.

L.Y. Funding acquisition, Resources,Writing -Review & Editing..

F.P. Funding acquisition, Resources,Writing -Review & Editing.

M.B., F.B., I.N., M.P., Funding acquisition, Resources.

A.M.B. Funding acquisition, Writing -Review & Editing.

P.B. Funding acquisition, Project administration, Resources, Data Curation, Writing -Review & Editing.

E.M. Funding acquisition, Project administration, Resources, Data Curation, Conceptualization, Methodology, Supervision, Writing -Original Draft.

## Acknowledgments

Training data were provided by the Human Connectome Project, WU-Minn Consortium (Principal Investigators: David Van Essen and Kamil Ugurbil; 1U54MH091657) funded by the 16 NIH Institutes and Centers that support the NIH Blueprint for Neuroscience Research; and by the McDonnell Center for Systems Neuroscience at Washington University.

Test data were provided by the non-funded StratiBip network initiative (Principal Investigator: Paolo Brambilla). The support from the StratiBip network members in collecting clinical and MRI data for the StratiBip dataset is acknowledged; for the Fondazione IRCCS Santa Lucia, we wish to acknowledge Daniela Vecchio for providing a key contribution to the data collection stage.

## Notes

### Competing Interest Statement

Authors declare no competing interests, except L.Y. who reports consultant/speaker fees from Alkermes, Allergan (currently Abbvie), Sumitomo Pharma, Intracellular Therapies, LivaNova, Merck, Sanofi, and Sunovion, and grants from Allergan (now AbbVie), CIHR, and Sumitomo, outside the submitted work, over the last 3 years.

