## Supplementary material for "A generalizable normative deep autoencoder for brain morphological anomaly detection: application to the multi-site StratiBip dataset on bipolar disorder in an external validation framework": Suplementary Information

*Corresponding author

^Equally contributing

***Indice***

***Suplementary Information***

*StratiBip dataset exclusion criteria*

*SPM/CAT-12 MRI preprocessing*

*ComBat model*

*Harmonization efficacy evaluation*

*Removing biocovariates effects via linear regression*

*Hyperparameter tunning*

***Suplementary Tables***

*Sup. Tab. 1- Comparison between HCP-YA, StratiBip-HC and StratiBip-BD*

*Sup. Tab. 2- StratiBip individual centers demographic information*

*Sup. Tab. 3- Pre vs. Post Harmonization distribution comparisons*

*Sup. Tab. 4 - SVM site classification pre. vs post harmonization by group and by feature set*

*Sup. Tab. 5- Significant associations between biocovariates and ROIs: pre vs. post harmonization*

*Sup. Tab. 6- Correlation between LR* $\beta$ *coefficients: pre. vs post harmonization*

*Sup. Tab. 7*- *Changes in significant associations between biocovariates and ROIs by diagnostic group: pre vs. post: harmonization*

*Sup. Tab. 8*- *Correlation between LR* $\beta$ *coefficients by diagnostic group: pre vs. post harmonization*

*Sup. Tab. 9*- *First hyperparameter search space*

*Sup. Tab. 10*- *Second hyperparameter search space*

*Sup. Tab. 11*- *Mass-univariate analysis on raw data HC vs. BD*

*Sup. Tab. 12*- *StratiBip individual sites MRI acquistion information*

*Sup. Tab. 13*- *DK40 cortical thickness region names*

*Sup. Tab. 14*- *CoBra Lab volumes region names*

***Suplementary Figures***

*Sup. Fig.1- Left: Age distributions; Right: Sex proportions*

*Sup. Fig.2- Boxplot mean feature distributions across sites and groups: pre vs. post harmonization*

*Sup. Fig.3- Dataset UMAP 2D projections : pre vs. post harmonization*

*Sup. Fig.4- AE model training evolution*

*Sup. Fig.5- 95% CI Features MDS*

*Sup. Fig.6-* *HCP-YA mZ scores features histograms with respective 99th percentile threshold*

*Sup. Fig.7-* *Features mZ distributions for HC and BD group*

**Suplementary Information | StratiBip dataset exclusion criteria**

All subjects were recruited and assessed at each individual site after signing a written and informed consent approved by the competent Ethical Committees, in accordance with the principles in the Declaration of Helsinki. The clinical assessment was performed by professional psychiatrists and diagnosis was based on the Diagnostic and Statistical Manual of Mental Disorders (DSM-IV) criteria for Axis I disorders (SCID-I) [27], with the exception of participants recruited in the Vancouver site (site id=7), for which diagnosis was based on the Mini International Neuropsychiatry Interview [28]. The participants exclusion criteria were: comorbidities, intellectual disability, pregnancy, history of epilepsy, major medical and neurological disorders, neuroleptic treatment in the last 3 months, drug or alcohol abuse in the last 6 months, and medical conditions affecting the immune system.

**Supplementary Information |** **SPM/CAT-12 MRI preprocessing**

The subjects’ T1-weighted images were segmented into brain tissues using SPM12 segmentation and ROI volumes processed with the CoBra atlases. We used the CAT12 pre-processing modules included in the standard voxel-based morphometry (VBM) starting with spatial registration using DARTEL (Diffeomorphic Anatomical Registration Through Exponentiated Lie algebra) resulting images were normalized to a reference template brain,t he Montreal Neurological Institute (MNI) space, followed by brain tissue segmentation, bias correction of intensity non-homogeneities, spatial normalization, modulation with the Jacobian determinant derived from the previous step to preserve the total amount of signal from each region, and spatial smoothing with a 3D Gaussian kernel with full-width half maximum of 6 mm. Surface-based morphometry was also performed to estimate cortical thickness using the projection-based thickness method. Each subject's total intracranial volume (TIV) was estimated. After a visual inspection of the tissue segmentation quality, we performed the CAT12-based quality check by verifying the spatial registration across images and identifying images with an intensity covariance below two standard deviations.

**Supplementary Information | ComBat model**

ComBat models biological covariates, such as age, sex, apriori, with a linear regression. The model starts by standardizing data to zero mean and unit variance, and removing from data the variance associated with the biological covariates, $X_{ji}^{T}\beta_{v}$, which are estimated at net of site by including them in the design matrix $X$ but not removing them during standardization, $y_{ijv}^{Standardized}=\frac{y_{ijv}- \hat{a}_{v}- X_{j}\hat{\beta}_{v}}{\hat{\sigma}_{v}}$, where $\hat{a}_{v}$ and $\hat{\sigma}_{v}$ are the standardization parameters, estimated mean, and variance across feature $v$. The standardized data is assumed to follow a normal distribution, $y_{ijv}^{Standardized}\sim N(\gamma_{iv},\delta_{iv})$. Then, the parametric model assumes a normal distribution for the additive site effect, $\gamma_{iv}\sim N\left( Yi,\tau^{2} \right),$and an inverse gamma for the multiplicative effect, $\delta_{iv}\sim IG(\lambda_{i},\theta_{i})$. ComBat parameter estimations ($\hat{a}_{v}$, $\hat{\sigma}_{v}$*,* $\gamma_{iv}$, $\delta_{iv}$) were applied in the StratiBip external test set in the following manner: first the estimated standardization statistics (based on HCP-YA reference) are used to standardize all test set data (HC and BD StratiBip). Second, before removing the estimated site-effects, biocovariates were accounted for, using the mean effects by averaging the estimated biocovariates coefficients, $\hat{\beta}_{v}$, across the reference site subjects (HCP-YA), $mod\_mean\left[ n_{subj}, n_{features} \right]=X_{i=HCP, j}{\times\hat{\beta}}_{v}$, thus ${mod\_mean}^{avg}=\frac{\sum_{j} {mod\_mean}_{i=HCP, j}}{{n\_subj}_{i=HCP}}$. The final data adjustment for all subjects in the external test sets was given by: $\hat{y}_{ijv}^{M-ComBat}=\frac{y_{ijv}-\hat{a}_{i=HCP,v}-{mod\_mean}^{avg}-\gamma_{iv}^{*}}{\delta_{iv}^{*}}+\hat{\alpha}_{i=HCP,v}+{mod\_mean}^{avg}$.

**Supplementary Information | Harmonization efficacy evaluation**

To evaluate the efficacy of site effects removal we first performed a thorough inter-site comparison between data distributions, before and after harmonization. Using a Kruskal Wallis ANOVA test we compared the mean CT, GMV, and WMV distribution between sites, separately on HC and BD data, before and after harmonization. For HC data, the 7 StratiBip sites and the HCP-YA were compared, followed by a Dunnett posthoc test in case of significant differences, using the HCP-YA as reference. On the other hand, for the BD group, only the 7 StratiBip sites were compared, as site differences between HCP-YA and BD could be confounded by group differences, followed by a TukeyHSD posthoc test in case of significant differences. Next, we assessed site-dependent clusterization on the entire dataset via UMAP non-linear dimensionality reduction method. UMAP embeds data by finding a low-dimensional data projection that has the closest possible equivalent fuzzy topological structure. This method was used to unveil the cluster-like structure, naturally present in data induced by the differences associated with site effects. Furthermore, we trained a support vector machine (SVM) classifier to predict site origin and assess the successful removal of site effects via decreasing model performance after data harmonization. First, all data was combined (N=1672) and split in 50% training (N=836) and 50% test set (N=836), stratifying for site. Then, with the model probability site-origin predictions we constructed the receiving operating characteristic curve (ROC) and evaluated model performance through the area under the curve (AUC-ROC) and f1-score. In a second analysis, we repeated the procedure by feature-type and by group. We used the *SVC()* function from *sklearn* to implement the SVM classifier with C=1, linear kernal, one-versus-rest decision function and a weighted class balance, and the *roc_auc_score* from *metrics* with a *one-versus-rest* multiclass strategy and a weighted average. To assert the total preservation of biological covariates effects, StratiBip intra-site linear regression (LR) models were computed before and after harmonization. Age, sex, and diagnosis were included as independent variables, and each brain feature was modelled as dependent variable, Eq. 3. This resulted in 170 models, computed with raw data and harmonized data, within each site (N=170x2x7). Mainly we assessed two conditions, i) the stability of biocovariates beta coefficients significance, before and after harmonization, and ii) the beta coefficients Spearman correlation, before and after harmonization. To satisfy the criteria of biological effects total preservation after M-ComBat harmonization, the significant associations between brain features and biocovariares have to be maintained after harmonization. The significance was assessed through the corrected p-values for multiple comparisons using the *holm-sidak* strategy, for an alpha level =0.05. Additionally, we expected high correlations between unharmonized and harmonized LR beta coefficients to consolidate the evidence for biocovariates effects preservation after harmonization. Finally, to show that M-ComBat harmonization performed similarly for HC and BD groups, despite all the models parameters being normatively estimated (i.e. exclusively in the HC group), we repeated the previous analysis by performing separate LR models for each group. All the models were computed using the *GLM()* function from *statsmodel* library using the Gaussian distribution family and identity link function.

$$y_{v:CT}^{pre\_harm}=\beta_{v0}+{\beta x}_{age}+{\beta x}_{sex}+\beta x_{diagn}$$

(3)

$$y_{v:volumes}^{pre\_harm}=\beta_{v0}+{\beta x}_{age}+{\beta x}_{sex}+\beta x_{diagn}+{\beta x}_{harm\_TIV}$$

**Supplementary information | Removing biocovariates effects via linear regression**

Each brain feature was modelled as the dependent variable, $Y_{v:[CT;volumes],i,j}^{ComBat}$, accounting for 170 independent models. The beta coefficients for age, sex, regarding CT features, and age, sex, and harmonized TIV regarding volumetric features were estimated in the HCP training set, $Y_{v:CT,volumes, i,j}^{harm}=\hat{const}_{v,i=HCP,j}+ \hat{\beta}_{v,i=HCP,j}X_{age}+ \hat{\beta}_{v,i=HCP,j}X_{sex}+ \hat{\beta}_{v=volumes,i=HCP,j}X_{harm\_TIV}$. Next, all the data is corrected for subject-specific biological covariates,$X_{j, cov}$, by removing its explained variance from respective brain feature: $Y_{vj}^{corr}=Y_{vj}^{harm}- \hat{\beta}_{v,i=HCP, j}X_{j,cov}$ , where $Y^{corr}$is residualized with respect to $X_{j,cov}$. Brain features were standardized using the statistics estimated on the HCP training set, before and after the LR stage, with *StandardScaler()* object from the *sklearn* python library.

**Supplementary Information | Hyperparameter tunning**

To find the most suitable network architecture we employed a 10-fold CV model hyperparameter tunning. We perform a first random search (RS) for 3 fixed batch sizes: 32, 64 and 124 with 700 iterations. Then, we inspected the best results and reduced the search space by keeping the best batch size, learning rates, epochs, and network layers dimensions that repeatedly performed better. With this reduced search space we re-perform a RS with 700 iterations by changing the random state seed. Additionally, an early stopping strategy was implemented with a tolerance of 100 epochs on the validation *MSE*. In each CV fold, we applied the CR pipeline before model training and validation, including the biocovariates’ LR and data standardization: all model coefficients were estimated on the 9 HCP-YA data partitions used as training set and then applied to all including the 10^th^ left-over partition used as validation set. Finally, the hyperparameter combination that resulted in a lower validation set average MSE across the 10-fold iterations was chosen.

**Supplementary Table 1 | Comparison between HCP-YA, StratiBip-HC and StratiBip-BD**

The age distributions between the three groups were significantly different according to a Kruskal-Wallis ANOVA test. The post-hoc pairwise comparison employing the Mann-Whitney U test with a Bonferroni correction for multiple comparisons showed that all groups differed among each other in terms of age distributions (HCvs.BD p=4.92e-6; HCvsHCP p=1.73e-6; HCPvsBD p=2.95e-2). On the other hand, based on a Chi-Square test of independence, no significant differences among the three groups were found for sex proportions (χ^2^(2, N=1659)=0.6338, p=0.728).

| **Group** | **Numerosity** | **Age**  Mean ± std \| median [25^th^;75^th^ ] | **Kruskal Wallis ANOVA** | **Post-hoc** | **Sex** | | **Chi-Square test** |
| --- | --- | --- | --- | --- | --- | --- | --- |
|  |  |  |  |  | **M** | **F** |  |
| HCP-YA | 1109 | 28.811 ±3.693  29.0 [25%: 26.0; 75%:32.0] | 34.85 p=2.702e10-8 | HCPvsHC p=1.73e-6  HCPvsBD p=2.95e-2 | 505 | 604 | χ^2^(0.6338 p=0.728) |
| HC-StratiBip | 363 | 27.741±3.999  27.0 [25%: 25.0 ; 75%: 31.0 ] |  | HCvsBD p=4.92e-6 | 174 | 189 |  |
| BD-StratiBip | 187 | 29.625 ±4.371  30.0 [25%: 26.0; 75%: 33.0] |  | - | 86 | 101 |  |

**Supplementary Table 2 | StratiBip individual centers demographic information**

Detailed demographic information for each StratiBip test set cohort is presented in this table. A 2x2 Chi-square test of independence was performed to investigate differences in sex distributions between the two groups, HC and BD, within each site. Instead, a Fisher Exact test was used for sites 1 and 3 due to low sample numerosity, signed with *. For comparing age distributions between the two groups, a Kruskall-Wallis ANOVA was used. When information is available only for a subset of subjects, the specific numerosity is presented *(n=X)*.

| **Site** | **Diagnosis** | | **Sex** | | **Χ^2^ pvalue** | **Age**  **(years)** | | **ANOVA pvalue** | **Education** | **BD-type** | | | **BD Age**  **Onset** |
| --- | --- | --- | --- | --- | --- | --- | --- | --- | --- | --- | --- | --- | --- |
|  |  |  |  |  |  |  |  |  |  | **BD-I** | | **BD-II** |  |
| **1-** | 74 | **HC** | **M** | 31 | 0.695* | 28.871± 3.827 | 28.162 +- 3.720 | 0.261 | 19.96± 1.44  *(n=72)* |  | | |  |
|  |  |  | **F** | 43 |  | 27.651± 3.598 |  |  |  |  |  |  |  |
|  | 3 | **BD** | **M** | 3 |  | 30.333± 2.310 | - |  | 14.67± 2.89 | 2 | 1 | | 22.67± 3.51 |
|  |  |  | **F** | - |  | - |  |  |  |  |  |  |  |
| **2-** | 93 | **HC** | **M** | 40 | 0.469 | 30.175± 4.050 | 29.227± 4.338 | 0.023 | 16.58± 2.63 |  | | |  |
|  |  |  | **F** | 53 |  | 27.262± 4.447 |  |  |  |  |  |  |  |
|  | 58 | **BD** | **M** | 30 |  | 30.765± 3.617 | 30.895± 3.946 |  | 14.51± 2.85 | 31 | 27 | | 20.75± 5.02  *(n=57)* |
|  |  |  | **F** | 28 |  | 31.036 ±4.333 |  |  |  |  |  |  |  |
| **3-** | 89 | **HC** | **M** | 47 | 1* | 26.106± 3.643 | 26.652± 3.940 | 0.085 | 16.71± 2.41  *(n=87)* |  | | |  |
|  |  |  | **F** | 42 |  | 27.262± 4.208 |  |  |  |  |  |  |  |
|  | 13 | **BD** | **M** | 8 |  | 29.625± 5.423 | 29.154± 5.064 |  | 14.77± 3.63 | 12 | 1 | | - |
|  |  |  | **F** | 5 |  | 28.400± 4.960 |  |  |  |  |  |  |  |
| **4-** | 23 | **HC** | **M** | 16 | 0.496 | 26.875± 3.722 | 26.694± 3.509 | 0.254 | - |  | | |  |
|  |  |  | **F** | 7 |  | 26.286± 3.200 |  |  |  |  |  |  |  |
|  | 6 | **BD** | **M** | 5 |  | 27.800± 3.114 | 27.838± 2.787 |  | 15.50± 2.74 | 3 | 3 | | 19.67± 6.25 |
|  |  |  | **F** | 1 |  | 28 |  |  |  |  |  |  |  |
| **5-** | 44 | **HC** | **M** | 24 | 0.906 | 26.350± 3.345 | 27.295± 3.867 | 1.2e-5 | - |  | | |  |
|  |  |  | **F** | 20 |  | 28.083± 4.159 |  |  |  |  |  |  |  |
|  | 37 | **BD** | **M** | 13 |  | 31.462± 4.539 | 31.837± 4.168 |  | 12.89± 3.59  *(n=36)* | 20 | 14 | | 24.42± 5.60  *(n=36)* |
|  |  |  | **F** | 24 |  | 32.042± 4.038 |  |  |  |  |  |  |  |
| **6-** | 25 | **HC** | **M** | 11 | 0.855 | 27.909± 2.700 | 27.920± 3.054 | 0.405 | - |  | | |  |
|  |  |  | **F** | 14 |  | 27.929± 3.407 |  |  |  |  |  |  |  |
|  | 40 | **BD** | **M** | 16 |  | 30.437± 3.669 | 28.800± 3.969 |  | - | 40 | - | | 18.15± 5.87 |
|  |  |  | **F** | 24 |  | 27.708± 3.850 |  |  |  |  |  |  |  |
| **7-** | 15 | **HC** | **M** | 5 | 0.505 | 26.200± 3.493 | 25.533± 2.850 | 0.818 | - |  | | |  |
|  |  |  | **F** | 10 |  | 25.200± 2.616 |  |  |  |  |  |  |  |
|  | 30 | **BD** | **M** | 11 |  | 25.364± 3.295 | 26.033± 3.508 |  | - | 30 | - | | 21.73± 5.70 |
|  |  |  | **F** | 19 |  | 26.421± 3.656 |  |  |  |  |  |  |  |

**Supplementary Table 3 | Pre vs. Post Harmonization distribution comparisons**

HC raw data distributions were compared among the eight sites (7 StratiBip cohorts and HCP-YA), labelled as *all_HC*, for each feature type (CT, GMV, WMV) using a Kruskall-wallis ANOVA hypothesis testing. Afterwards, if statistically significant differences were present, a post-hoc Dunnet test was performed using HCP-YA as reference. For the StratiBip-BD group, the feature set distributions were compared among the 7 StratiBip sites and a tukeyHSD posthoc test was used as follow-up for pairwise comparisons if significant differences were found. Post-hoc pvalues results were corrected for multiple comparisons using the Holm-sidak correction, due to a number of comparisons equal or higher than 7.

| **Feature Type Distribution for two groups** | **Pre-Harmonization**  **p<0.05 uncorrected** | **Post-hoc Holm-sidak correction**  **p<0.05** | **Post-Harmonization**  **p<0.05 uncorrected** | **Post-hoc**  **Holm-sidak correction**  **p<0.05** |
| --- | --- | --- | --- | --- |
| **CT_all_HC_** | p=9.53e-82 | All* except site 6 and site 7 | p=0.680 | No differences |
| **CT_BD_** | p=3.108e-21 | Only site 5 with 2,3,4,6,7 and site 2 with 7 | **p=0.000975** | Differences between sites 4 and 6 (p=0.013) |
| **GMV_all_HC_** | p=1.148e-94 | All | p=0.913 | No differences |
| **GMV_BD_** | p=9.11e-13 | Only site 5 with 2,3,4,6,7 | **p=0.018** | No differences survived multiple comparison correction |
| **WMV_all_HC_** | p=2.68e-29 | All* except site 4 | p=0.926 | No differences |
| **WMV_BD_** | p=4.918e-12 | Only site 5 with 1,2,3,6,7 | p=0.0721 | No differences |

*All pairwise comparisons revealed statistically significant differences on the distributions except for (…).

**Supplementary Table 4 | SVM site classification pre. vs post harmonization by group and by feature set**

The SVM site classification performance was inspected before and after data harmonization by group and by feature set, through the weighted average AUC-ROC and f1-score. The ability to predict site origin was hampered after data harmonization, as seen by the decrease in AUC-ROC, evidence for the successful removal of the encoded site effects in the neuroimaging-derived features.

| SVM model performance | | AUC-ROC \| F1-Score | | | | |
| --- | --- | --- | --- | --- | --- | --- |
|  |  | All features  (n=170) | CT  (n=68) | | GMV  (n=50) | WMV  (n=52) |
| All data  (N_train_=829 \| N_test_=830) | Raw Data | 0.99 \| 0.95 | 0.99 \| 0.92 | | 0.99 \| 0.88 | 0.99 \| 0.93 |
|  | Harmonized | 0.57 \| 0.23 | 0.55 \| 0.14 | | 0.54 \| 0.13 | 0.52 \| 0.10 |
| HC-StratiBip data  (N_train_=181 \| N_test_=182) | Raw Data | 0.99 \| 0.88 | 0.96 \| 0.79 | | 0.91 \| 0.62 | 0.96 \| 0.70 |
|  | Harmonized | 0.74 \| 0.06 | 0.60 \| 0.04 | | 0.62 \| 0.08 | 0.61\| 0.11 |
| BD-StratiBip data  (N_train_=93 \| N_test_=94) | Raw Data | 0.96 \| 0.79 | 0.95 \| 0.81 | 0.82 \| 0.56 | | 0.94 \| 0.72 |
|  | Harmonized | 0.70 \| 0.43 | 0.72 \| 0.40 | 0.62 \| 0.30 | | 0.62 \| 0.25 |

**Supplementary Table 5 | Significant associations between biocovariates and ROIs: pre vs. post harmonization**

To evaluate the successfulness of the M-ComBat harmonization, we assessed the preservation of biological covariates. Independent linear regression (LR) models were computed separately for data of each site, before and after harmonization, using age, sex, and diagnosis as independent variables for the 68 CT brain features, whereas harmonized TIV was added as an independent variable for GMV and WMV features, , $Y_{v:CT,volumes, i,j}^{pre/post\_harm}=\hat{const}_{v,i,j}+ \hat{\beta}_{v,i,j}X_{age}+ \hat{\beta}_{v,i,j}X_{sex}+ \hat{\beta}_{v=volumes,i,j}X_{harm\_TIV}$, consistent with the M-ComBat pipeline. We report the number of identified significant associations between biological covariates and brain features, i.e., the biocovariates for which $\hat{\beta}_{v,i,j}$ presented a significant pvalue (p<0.05), pre and post-harmonization, after correcting for multiple comparisons using the Holm-sidak strategy. The reported results indicate that the biological covariates effects were totally preserved in the harmonized data as there were no changes in their significance after this process.

| Features | Sites: | Age | Sex | Diagnosis | TIV |
| --- | --- | --- | --- | --- | --- |
|  |  | **1 \| 2 \| 3 \| 4 \| 5 \| 6 \| 7** | **1 \| 2 \| 3 \| 4 \| 5 \| 6 \| 7** | **1 \| 2 \| 3 \| 4 \| 5 \| 6 \| 7** | **1 \| 2 \| 3 \| 4 \| 5 \| 6 \| 7** |
| CT | Pre | 0 \| 0 \| 0 \| 0 \| 0 \| 1 \| 10 | 0 \| 0 \| 0 \| 0 \| 0 \| 0 \| 0 | 0 \| 0 \| 0 \| 1 \| 0 \| 0 \| 0 | - |
|  | Post | 0 \| 0 \| 0 \| 0 \| 0 \| 1 \| 10 | 0 \| 0 \| 0 \| 0 \| 0 \| 0 \| 0 | 0 \| 0 \| 0 \| 1 \| 0 \| 0 \| 0 | - |
| GMV | Pre | 0 \| 0 \| 0 \| 0 \| 0 \| 0 \| 0 | 0 \| 0 \| 0 \| 0 \| 0 \| 0 \| 1 | 4 \| 0 \| 0 \| 0\| 0 \| 0 \| 0 | 18 \| 44 \| 46 \| 25 \| 45 \| 42 \| 23 |
|  | Post | 0 \| 0 \| 0 \| 0 \| 0 \| 0 \| 0 | 0 \| 0 \| 0 \| 0 \| 0 \| 0 \| 1 | 4 \| 0 \| 0 \| 0 \| 0 \| 0 \| 0 | 18 \| 44 \| 46 \| 25 \| 45 \| 42 \| 23 |
| WMV | Pre | 0 \| 0 \| 0 \| 0 \| 0 \| 0 \| 0 | 0 \| 0 \| 1 \| 0 \| 0 \| 0 \| 0 | 0 \| 0 \| 1 \| 0 \| 0 \| 0 \| 0 | 27 \| 42 \| 40 \| 24 \| 36 \| 47 \| 11 |
|  | Post | 0 \| 0 \| 0 \| 0 \| 0 \| 0 \| 0 | 0 \| 0 \| 1 \| 0 \| 0 \| 0 \| 0 | 0 \| 0 \| 1 \| 0 \| 0 \| 0 \| 0 | 27 \| 42 \| 40 \| 24 \| 36 \| 47 \| 11 |
| TIV | Pre | 0 \| 0 \| 0 \| 0 \| 0 \| 0 \| 0 | 1 \| 0 \| 0 \| 0 \| 0 \| 0 \| 1 | 1 \| 0 \| 0 \| 0 \| 0 \| 0 \| 0 | - |
|  | Post | 0 \| 0 \| 0 \| 0 \| 0 \| 0 \| 0 | 1 \| 0 \| 0 \| 0 \| 0 \| 0 \| 1 | 1 \| 0 \| 0 \| 0 \| 0 \| 0 \| 0 | - |

**Supplementary Table 6 | Correlation between LR** $\beta$ **coefficients: pre vs. post harmonization**

We computed the Spearman correlation between biocovariates beta coefficients, $\beta,$ pre and post-harmonization, from the intra-site LR models. All the correlations resulted statistically significant (p<10^-22^) with r coefficients > 0.96. The stability of significance in intra-site models between pre and post-harmonization, and the high beta coefficient correlations reported, ensures that the harmonization pipeline effectively preserved the biological covariates in the StratiBip external sets. Importantly, potential diagnostic effects in data were also preserve as seen from the
$\beta_{diagnosis}$ stability, i.e. high significant correlation, between pre. vs. post harmonization models.

| Features | Spearman correlation r ($\beta_{raw}, \beta_{harm})$ | | | | | | | |
| --- | --- | --- | --- | --- | --- | --- | --- | --- |
|  | Sites: | 1 | 2 | 3 | 4 | 5 | 6 | 7 |
| CT | $\beta_{age}$ | 0.98 | 0.99 | 0.99 | 0.99 | 0.97 | 0.99 | 0.99 |
|  | $\beta_{sex}$ | 0.98 | 0.99 | 0.99 | 0.99 | 0.99 | 0.99 | 0.99 |
|  | $\beta_{diagnosis}$ | 0.97 | 0.99 | 0.99 | 0.97 | 0.99 | 0.97 | 0.99 |
| GMV | $\beta_{age}$ | 0.99 | 0.99 | 0.99 | 0.99 | 0.99 | 0.99 | 0.99 |
|  | $\beta_{sex}$ | 0.99 | 0.99 | 0.99 | 0.99 | 0.99 | 0.99 | 0.99 |
|  | $\beta_{diagnosis}$ | 0.99 | 0.99 | 0.99 | 0.99 | 0.99 | 0.98 | 0.99 |
|  | $\beta_{TIV}$ | 0.99 | 0.99 | 0.99 | 0.99 | 0.98 | 0.99 | 0.99 |
| WMV | $\beta_{age}$ | 0.99 | 0.99 | 0.99 | 0.99 | 0.99 | 0.98 | 0.99 |
|  | $\beta_{sex}$ | 0.99 | 0.99 | 0.98 | 0.99 | 0.99 | 0.99 | 0.99 |
|  | $\beta_{diagnosis}$ | 0.99 | 0.99 | 0.99 | 0.99 | 0.99 | 0.99 | 0.99 |
|  | $\beta_{TIV}$ | 0.96 | 0.97 | 0.97 | 0.97 | 0.98 | 0.98 | 0.99 |

**Supplementary Table 7 | Changes in significant associations between biocovariates and ROIs by diagnostic group: pre vs. post harmonization**

To ascertain equal performance in terms of biological covariates preservation for both HC and BD groups, we performed an intra-site LR analysis, identical to the one presented in Sup. Table 5, separately for the two groups. In this case, diagnosis was not modeled within the LR. The changes in significance were extracted by observing the changes in pvalue for each $\beta$ in the intra-site LR’s. The analysis on data from site 1 was not performed, as this site contained only 3 BD subjects. Similarly, as in the previous analysis, there were no changes in significant associations for both HC and BD groups.

| Δ significance | Group | Age | Sex | TIV |
| --- | --- | --- | --- | --- |
|  |  | **Sites 1 \| 2 \| 3 \| 4 \| 5 \| 6 \| 7** | **1 \| 2 \| 3 \| 4 \| 5 \| 6 \| 7** | **1 \| 2 \| 3 \| 4 \| 5 \| 6 \| 7** |
| Cortical Thickness  (N=68) | HC | 0 \| 0 \| 0 \| 0 \| 0 \| 0 \| 0 | 0 \| 0 \| 0 \| 0 \| 0 \| 0 \| 0 | - |
|  | BD | -*\| 0 \| 0 \| 0 \| 0 \| 0 \| 0 | -*\| 0 \| 0 \| 0 \| 0 \| 0 \| 0 | - |
| Gray Matter Volumes  (N=50) | HC | 0 \| 0 \| 0 \| 0 \| 0 \| 0 \| 0 | 0 \| 0 \| 0 \| 0 \| 0 \| 0 \| 0 | 0 \| 0 \| 0 \| 0 \| 0 \| 0 \| 0 |
|  | BD | -*\| 0 \| 0 \| 0 \| 0 \| 0 \| 0 | - *\|0 \| 0 \| 0 \| 0 \| 0 \| 0 | -*\| 0 \| 0 \| 0 \| 0 \| 0 \| 0 |
| White Mattern Volumes  (N=52) | HC | 0 \| 0 \| 0 \| 0 \| 0 \| 0 \| 0 | 0 \| 0 \| 0 \| 0 \| 0 \| 0 \| 0 | 0 \| 0 \| 0 \| 0 \| 0 \| 0 \| 0 |
|  | BD | -*\| 0 \| 0 \| 0 \| 0 \| 0 \| 0 | -*\| 0 \| 0 \| 0 \| 0 \| 0 \| 0 | -*\| 0 \| 0 \| 0 \| 0 \| 0 \| 0 |
| Total Intracranial Volume | HC | 0 \| 0 \| 0 \| 0 \| 0 \| 0 \| 0 | 0 \| 0 \| 0 \| 0 \| 0 \| 0 \| 0 | - |
|  | BD | -*\| 0 \| 0 \| 0 \| 0 \| 0 \| 0 | -*\| 0 \| 0 \| 0 \| 0 \| 0 \| 0 | - |

*only 3 subjects available;

**Supplementary Table 8 | Correlation between LR** $\beta$ **coefficients by diagnostic group: pre vs. post harmonization**

We computed the Spearman correlation between biocovariates beta coefficients, $\beta,$ pre and post-harmonization, from the intra-site by diagnostic group LR models. All the correlations resulted statistically significant (p<10^-31^) with r coefficients > 0.96. The stability of significance in intra-site by group LR models between pre and post-harmonization, and the high beta coefficient correlations reported, ensures that the harmonization pipeline effectively preserved the biological covariates similarly in both groups in the StratiBip external sets.

| Group | Features | Spearman correlation r ($\beta_{raw}, \beta_{harm})$ | | | | | | | |
| --- | --- | --- | --- | --- | --- | --- | --- | --- | --- |
|  |  | Sites: | 1 | 2 | 3 | 4 | 5 | 6 | 7 |
| HC | CT | $\beta_{age}$ | 0.98 | 0.99 | 0.98 | 0.99 | 0.98 | 0.97 | 0.99 |
|  |  | $\beta_{sex}$ | 0.99 | 0.99 | 0.99 | 0.99 | 0.99 | 0.99 | 0.99 |
|  | GMV | $\beta_{age}$ | 0.99 | 0.99 | 0.99 | 0.99 | 0.99 | 0.99 | 0.99 |
|  |  | $\beta_{sex}$ | 0.98 | 0.99 | 0.98 | 0.99 | 0.99 | 0.98 | 0.99 |
|  |  | $\beta_{TIV}$ | **0.96** | 0.98 | 0.98 | **0.97** | 0.98 | 0.98 | 0.99 |
|  | WMV | $\beta_{age}$ | 0.99 | 0.99 | 0.99 | 0.99 | 0.99 | 0.98 | 0.99 |
|  |  | $\beta_{sex}$ | 0.98 | 0.99 | 0.98 | 0.99 | 0.99 | 0.98 | 0.99 |
|  |  | $\beta_{TIV}$ | 0.96 | 0.98 | 0.98 | 0.97 | 0.98 | 0.98 | 0.99 |
| BD | CT | $\beta_{age}$ | - | 0.99 | 0.99 | 0.98 | 0.99 | 0.99 | 0.98 |
|  |  | $\beta_{sex}$ | - | 0.99 | 0.99 | 0.99 | 0.99 | 0.99 | 0.99 |
|  | GMV | $\beta_{age}$ | - | 0.99 | 0.99 | 0.99 | 0.99 | 0.99 | 0.99 |
|  |  | $\beta_{sex}$ | - | 0.97 | 0.99 | 0.99 | 0.99 | 0.99 | 0.99 |
|  |  | $\beta_{TIV}$ | - | 0.98 | 0.99 | 0.99 | 0.98 | 0.99 | 0.98 |
|  | WMV | $\beta_{age}$ | - | 0.99 | 0.99 | 0.99 | 0.98 | 0.98 | 0.99 |
|  |  | $\beta_{sex}$ | - | 0.99 | 0.98 | 0.99 | 0.99 | 0.99 | 0.99 |
|  |  | $\beta_{TIV}$ | - | 0.97 | 0.98 | 0.99 | 0.98 | 0.99 | 0.99 |

**Supplementary Table 9 | First hyperparameter search space**

Batch size: **32**, 64, 124

Seed=42

| Network layers and dimensions | [150,120,80]; [150,100,80] ; [150,90,70] ; **[120,90,70] ; [120,100,80] ; [100,70] ; [120,80] ; [90] ; [100]** |
| --- | --- |
| Network bottleneck dimension | 20,25,30,35,40,45,50,55,60 |
| Learning Rate | **0.00001,0.000001,0.0001,0.005,0.00005,0.001, 0.0005**, lr_schedule (initial=0.05, exp_decay=0.9977), lr_schedule (initial=0.01, exp_decay= 0.9977) |
| Epochs | 500,**2000,1000** |
| Kernel L2 Regularizer | **0.001,0.0001,0.00001,0.01,0.005,**  **0.0005,0.1,0.00005** |

*****in bold the options that presented better results across all iterations of the RS. The other hyperparameters were discarded in the second round of RS.

**Supplementary Table 10 | Second hyperparameter search space**

The identified best hyperparameter combination was: first encoding layer= 100, latent dimension= 60, learning rate= 0.00005, batch size= 32, epochs= 2000, l2 kernel regularizer= 0.0005.

| Batch size= 32 | Seed= 123 |
| --- | --- |
| Network layers and dimensions | [120,90,70] ; [120,100,80], [100,70], [120,80], **[100]**, [80] |
| Network bottleneck dimension | 20,25,30,35,40,45,50,55,**60** |
| Learning Rate | 0.00001, 0.000001, 0.0001,0.001, 0.005, **0.00005**, 0.0005 |
| Epochs | **2000**,1000,3000,4000 |
| Kernel L2 Regularizer | 0.001,0.0001,0.00001,0.005, **0.0005**, 0.00005 |

**Supplementary Table 11 | Mass-univariate analysis on raw data HC vs. BD**

One feature was identified both in the mass-univariate analysis and AE-normative based model as being significantly different between HC and BD, the left WM volume adjacent to globus pallidus (green highlight), although hippocampus subregions CA4 and CA2_3 were common denominators in both findings. The overlap coefficient between the AE-normative based findings and comparable mass-univariate analysis was 0.14. Using a Benjamin-Hochenber FDR correction for multiple comparisons for a critical level=0.05, the identified features decreased from 21 to 4: bilateral bankssts and bilateral WM adjacent to globus pallidus, highlighted in table with bold font.

| MUA 1  p<0.05 | Cliffs Delta |
| --- | --- |
| **lbankssts** | **-1** |
| **rbankssts** | **0.191** |
| Lentorhinal | 0.117 |
| Rmiddletemporal | 0.138 |
| Lparahippocampal | 0.109 |
| Rparsorbitalis | 0.104 |
| Rparstriangularis | 0.112 |
| Ltemporalpole | 0.145 |
| GM lGloPal | -0.147 |
| GM rGloPal | -0.140 |
| **WM adjacent to lGloPal** | **0.179** |
| WM lAntCerebLI_II | 0.107 |
| WM adjacent to lAmy | -0.121 |
| WM lFor | 0.111 |
| WM lCA4 | -0.116 |
| WM lCA2_3 | 0.107 |
| WM lFimbra | 0.132 |
| **WM adjacent to rGloPal** | **0.177** |
| WM adjacent to rAmy | -0.120 |
| WM rFor | 0.141 |
| WM rCA4 | -0.114 |

**Suplementary Table 12 |** **StratiBip individual sites MRI acquistion information**

| **ID** | **Reference PI** | **Scanner** | **Sequence** | **Matrix Size** | **Voxel Size (m^3^)** |
| --- | --- | --- | --- | --- | --- |
| **1-AUOV-Verona Italy** | Marcella Bellani and Paolo Brambilla | Magnetom Allegra Syngo (Siemens, Erlangen, Germany) | T1-MPRAGE | 256x256x160 | 1.00x1.00.x1.00 |
| **2-FSL_ROME - Fondazione IRCCS Santa Lucia, Roma, Italy** | Fabrizio Piras | Philips Achieva 3T (Philips, Best, the Netherlands) | T13D-MPRAGE | 432x432x190 | 0.542x 0.524.x 0.900 |
| **3-JUH-University of Jena, Germany** | Igor Nenadic | Siemens Tim Trio (Siemens, Erlangen, Germany) | T1 Magnetization Prepared Rapid Gradient Echo (MP-RAGE) | 256x256x192 | 1.00x1.00.x1.00 |
| **4-MI-Milano Policlinico, Italy** | Paolo Brambilla | Philips Achieva 3T (Philips, Best, the Netherlands) | T1-Turbo Field Echo (TFE) 3D | 240x240x165 | 1.1x1.05x1.05 |
| **5-OSR-Ospedale San Raffaele, Milan, Italy** | Francesco Benedetti | Philips Intera (Philips, Best, the Netherlands) | T1-Fast Field Echo (FFE) | 256x256x220 | 0.9x0.9x0.8 |
| **6- PITTS-Pittsburgh, US** | Mary Philips | 3T Siemens Tim Trio | **-** | 192x256x192 | 1.00x1.00x1.00 |
| **7-UBC- University of British Columbia, Vancouver, Canada** | Lakshmi Yatham | Philips Achieva (Philips, Best, the Netherlands) | 3D TFE | 256x256x180 | 1.00x1.00x1.00 |

**Supplementary Table 13 | DK40 cortical thickness region names**

| **ID** | **Abbreviation** | **ROI Name** | **Lobe** |
| --- | --- | --- | --- |
| 2647065 | 'lbankssts' | Banks superior temporal sulcus | Temporal |
| 2647065 | 'rbankssts' |  |  |
| 10511485 | 'lcaudalanteriorcingulate' | Caudal anterior-cingulate cortex | Frontal |
| 10511485 | 'rcaudalanteriorcingulate' |  |  |
| 6500 | 'lcaudalmiddlefrontal' | Caudal middle frontal gyrus | Frontal |
| 6500 | 'rcaudalmiddlefrontal' |  |  |
| 3294840 | 'lcorpuscallosum' | Corpus Callosum | WM |
| 3294840 | 'rcorpuscallosum' |  |  |
| 6558940 | 'lcuneus' | Cuneus cortex | Occipital |
| 6558940 | 'rcuneus' |  |  |
| 660700 | 'lentorhinal' | Entorhinal cortex | Temporal |
| 660700 | 'rentorhinal' |  |  |
| 9231540 | 'lfusiform' | Fusiform gyrus | Temporal |
| 9231540 | 'rfusiform' |  |  |
| 14433500 | 'linferiorparietal' | Inferior parietal cortex | Parietal |
| 14433500 | 'rinferiorparietal' |  |  |
| 7874740 | 'linferiortemporal' | Inferior temporal gyrus | Temporal |
| 7874740 | 'rinferiortemporal' |  |  |
| 9180300 | 'listhmuscingulate' | Isthmus – cingulate cortex | Parietal |
| 9180300 | 'risthmuscingulate' |  |  |
| 9182740 | 'llateraloccipital' | Lateral occipital cortex | Occipital |
| 9182740 | 'rlateraloccipital' |  |  |
| 3296035 | 'llateralorbitofrontal' | Lateral orbital frontal cortex | Frontal |
| 3296035 | 'rlateralorbitofrontal' |  |  |
| 9211105 | 'llingual' | Lingual gyrus | Occipital |
| 9211105 | 'rlingual' |  |  |
| 4924360 | 'lmedialorbitofrontal' | Medial orbital frontal cortex | Frontal |
| 4924360 | 'rmedialorbitofrontal' |  |  |
| 3302560 | 'lmiddletemporal' | Middle temporal gyrus | Temporal |
| 3302560 | 'rmiddletemporal' |  |  |
| 3988500 | 'lparahippocampal' | Parahippocampal gyrus | Temporal |
| 3988500 | 'rparahippocampal' |  |  |
| 3988540 | 'lparacentral' | Paracentral lobule | Frontal |
| 3988540 | 'rparacentral' |  |  |
| 9221340 | 'lparsopercularis' | Pars opercularis | Frontal |
| 9221340 | 'rparsopercularis' |  |  |
| 3302420 | 'lparsorbitalis' | Pars orbitalis | Frontal |
| 3302420 | 'rparsorbitalis' |  |  |
| 1326300 | 'lparstriangularis' | Pars triangularis | Frontal |
| 1326300 | 'rparstriangularis' |  |  |
| 3957880 | 'lpericalcarine' | Pericalcarine cortex | Occipital |
| 3957880 | 'rpericalcarine' |  |  |
| 1316060 | 'lpostcentral' | Postcentral gyrus | Parietal |
| 1316060 | 'rpostcentral' |  |  |
| 14464220 | 'lposteriorcingulate' | Posterior-cingulate cortex | Parietal |
| 14464220 | 'rposteriorcingulate' |  |  |
| 14423100 | 'lprecentral' | Precentral gyrus | Frontal |
| 14423100 | 'rprecentral' |  |  |
| 11832480 | 'lprecuneus' | Precuneus cortex | Parietal |
| 11832480 | 'rprecuneus' |  |  |
| 9180240 | 'lrostralanteriorcingulate' | Rostral anterior cingulate cortex | Frontal |
| 9180240 | 'rrostralanteriorcingulate' |  |  |
| 8204875 | 'lrostralmiddlefrontal' | Rostral middle frontal gyrus | Frontal |
| 8204875 | 'rrostralmiddlefrontal' |  |  |
| 10542100 | 'lsuperiorfrontal' | Superior frontal gyrus | Frontal |
| 10542100 | 'rsuperiorfrontal' |  |  |
| 9221140 | 'lsuperiorparietal' | Superior parietal cortex | Parietal |
| 9221140 | 'rsuperiorparietal' |  |  |
| 14474380 | 'lsuperiortemporal' | Superior temporal gyrus | Temporal |
| 14474380 | 'rsuperiortemporal' |  |  |
| 1351760 | 'lsupramarginal' | Supramarginal gyrus | Parietal |
| 1351760 | 'rsupramarginal' |  |  |
| 6553700 | 'lfrontalpole' | Frontal pole | Frontal |
| 6553700 | 'rfrontalpole' |  |  |
| 11146310 | 'ltemporalpole' | Temporal pole | Temporal |
| 11146310 | 'rtemporalpole' |  |  |
| 13145750 | 'ltransversetemporal' | Transverse temporal cortex | Temporal |
| 13145750 | 'rtransversetemporal' |  |  |
| 2146559 | 'linsula' | Insula | Insula |
| 2146559 | 'rinsula' |  |  |

**Supplementary Table 14 | CoBra Atlas GM volumes region names**

| ROIid | ROIabbr | ROIname |
| --- | --- | --- |
| 1 | lStriatum | Left Striatum |
| 2 | lGloPal | Left Globus Pallidus |
| 3 | lTha | Left Thalamus |
| 11 | lAntCerebLI_II | Left Anterior Cerebellar Lobule I-II |
| 12 | lAntCerebLIII | Left Anterior Cerebellar Lobule III |
| 13 | lAntCerebLIV | Left Anterior Cerebellar Lobule IV |
| 14 | lAntCerebLV | Left Anterior Cerebellar Lobule V |
| 15 | lSupPostCerebLVI | Left Superior Posterior Cerebellar Lobule VI |
| 16 | lSupPostCerebCI | Left Superior Posterior Cerebellar Lobule Crus I |
| 17 | lSupPostCerebCII | Left Superior Posterior Cerebellar Lobule Crus II |
| 18 | lSupPostCerebLVIIB | Left Superior Posterior Cerebellar Lobule VIIB |
| 19 | lInfPostCerebLVIIIA | Left Inferior Posterior Cerebellar Lobule VIIIA |
| 20 | lInfPostCerebLVIIIB | Left Inferior Posterior Cerebellar Lobule VIIIB |
| 21 | lInfPostCerebLIX | Left Inferior Posterior Cerebellar Lobule IX |
| 22 | lInfPostCerebLX | Left Inferior Posterior Cerebellar Lobule X |
| 26 | lAmy | Left Amygdala |
| 31 | lHCA1 | Left Hippocampus CA1 |
| 32 | lSub | Left Subiculum |
| 33 | lFor | Adjacent to Left Fornix |
| 34 | lCA4 | Left CA4/Dentate Gyrus |
| 35 | lCA2_3 | Left CA2/CA3 |
| 36 | lStratum | Left Stratum Radiatum/Lacunosum/Moleculare |
| 37 | lFimbra | Adjacent to Left Fimbria |
| 38 | lMamBody | Adjacent to Left Mammillary body |
| 39 | lAlveus | Adjacent to Left Alveus |
| 101 | rStriatum | Right Striatum |
| 102 | rGloPal | Right Globus Pallidus |
| 103 | rTha | Right Thalamus |
| 111 | rAntCerebLI_II | Right Anterior Cerebellar Lobule I-II |
| 112 | rAntCerebLIII | Right Anterior Cerebellar Lobule III |
| 113 | rAntCerebLIV | Right Anterior Cerebellar Lobule IV |
| 114 | rAntCerebLV | Right Anterior Cerebellar Lobule V |
| 115 | rSupPostCerebLVI | Right Superior Posterior Cerebellar Lobule VI |
| 116 | rSupPostCerebCI | Right Superior Posterior Cerebellar Lobule Crus I |
| 117 | rSupPostCerebCII | Right Superior Posterior Cerebellar Lobule Crus II |
| 118 | rSupPostCerebLVIIB | Right Superior Posterior Cerebellar Lobule VIIB |
| 119 | rInfPostCerebLVIIIA | Right Inferior Posterior Cerebellar Lobule VIIIA |
| 120 | rInfPostCerebLVIIIB | Right Inferior Posterior Cerebellar Lobule VIIIB |
| 121 | rInfPostCerebLIX | Right Inferior Posterior Cerebellar Lobule IX |
| 122 | rInfPostCerebLX | Right Inferior Posterior Cerebellar Lobule X |
| 126 | rAmy | Right Amygdala |
| 131 | rHCA1 | Right Hippocampus CA1 |
| 132 | rSub | Right Subiculum |
| 133 | rFor | Adjacent to Right Fornix |
| 134 | rCA4 | Right CA4/Dentate Gyrus |
| 135 | rCA2_3 | Right CA2/CA3 |
| 136 | rStratum | Right Stratum Radiatum/Lacunosum/Moleculare |
| 137 | rFimbra | Adjacent to Right Fimbria |
| 138 | rMamBody | Adjacent to Right Mammillary body |
| 139 | rAlveus | Adjacent to Right Alveus |

**Supplementary Table 15 | CoBra Atlas WM volumes region names**

| ROIid | ROIabbr | ROIname |
| --- | --- | --- |
| 1 | lStriatum | Adjacent to Left Striatum |
| 2 | lGloPal | Adjacent to Left Globus Pallidus |
| 3 | lTha | Adjacent to Left Thalamus |
| 11 | lAntCerebLI_II | Left Anterior Cerebellar Lobule I-II |
| 12 | lAntCerebLIII | Left Anterior Cerebellar Lobule III |
| 13 | lAntCerebLIV | Left Anterior Cerebellar Lobule IV |
| 14 | lAntCerebLV | Left Anterior Cerebellar Lobule V |
| 15 | lSupPostCerebLVI | Left Superior Posterior Cerebellar Lobule VI |
| 16 | lSupPostCerebCI | Left Superior Posterior Cerebellar Lobule Crus I |
| 17 | lSupPostCerebCII | Left Superior Posterior Cerebellar Lobule Crus II |
| 18 | lSupPostCerebLVIIB | Left Superior Posterior Cerebellar Lobule VIIB |
| 19 | lInfPostCerebLVIIIA | Left Inferior Posterior Cerebellar Lobule VIIIA |
| 20 | lInfPostCerebLVIIIB | Left Inferior Posterior Cerebellar Lobule VIIIB |
| 21 | lInfPostCerebLIX | Left Inferior Posterior Cerebellar Lobule IX |
| 22 | lInfPostCerebLX | Left Inferior Posterior Cerebellar Lobule X |
| 23 | lAntCerebWM | Left Cerebellar White Matter |
| 26 | lAmy | Adjacent to Left Amygdala |
| 31 | lHCA1 | Left Hippocampus CA1 |
| 32 | lSub | Left Subiculum |
| 33 | lFor | Left Fornix |
| 34 | lCA4 | Left CA4/Dentate Gyrus |
| 35 | lCA2_3 | Left CA2/CA3 |
| 36 | lStratum | Left Stratum Radiatum/Lacunosum/Moleculare |
| 37 | lFimbra | Left Fimbria |
| 38 | lMamBody | Left Mammillary body |
| 39 | lAlveus | Left Alveus |
| 101 | rStriatum | Adjacent to Right Striatum |
| 102 | rGloPal | Adjacent to Right Globus Pallidus |
| 103 | rTha | Adjacent to Right Thalamus |
| 111 | rAntCerebLI_II | Right Anterior Cerebellar Lobule I-II |
| 112 | rAntCerebLIII | Right Anterior Cerebellar Lobule III |
| 113 | rAntCerebLIV | Right Anterior Cerebellar Lobule IV |
| 114 | rAntCerebLV | Right Anterior Cerebellar Lobule V |
| 115 | rSupPostCerebLVI | Right Superior Posterior Cerebellar Lobule VI |
| 116 | rSupPostCerebCI | Right Superior Posterior Cerebellar Lobule Crus I |
| 117 | rSupPostCerebCII | Right Superior Posterior Cerebellar Lobule Crus II |
| 118 | rSupPostCerebLVIIB | Right Superior Posterior Cerebellar Lobule VIIB |
| 119 | rInfPostCerebLVIIIA | Right Inferior Posterior Cerebellar Lobule VIIIA |
| 120 | rInfPostCerebLVIIIB | Right Inferior Posterior Cerebellar Lobule VIIIB |
| 121 | rInfPostCerebLIX | Right Inferior Posterior Cerebellar Lobule IX |
| 122 | rInfPostCerebLX | Right Inferior Posterior Cerebellar Lobule X |
| 123 | rAntCerebWM | Right Cerebellar White Matter |
| 126 | rAmy | Adjacent to Right Amygdala |
| 131 | rHCA1 | Right Hippocampus CA1 |
| 132 | rSub | Right Subiculum |
| 133 | rFor | Right Fornix |
| 134 | rCA4 | Right CA4/Dentate Gyrus |
| 135 | rCA2_3 | Right CA2/CA3 |
| 136 | rStratum | Right Stratum Radiatum/Lacunosum/Moleculare |
| 137 | rFimbra | Right Fimbria |
| 138 | rMamBody | Right Mammillary body |
| 139 | rAlveus | Right Alveus |

**Supplementary Figure 1 | Left: Age distributions; Right: Sex proportions**

Age distributions were statistically significantly different between the three groups. A post-hoc two-sided MWU test with a Bonferroni correction revealed all pairwise comparisons to be significantly different (HCvs.BD p=4.92e-6; HCvsHCP p=1.73e-6; HCPvsBD p=2.95e-2).

| 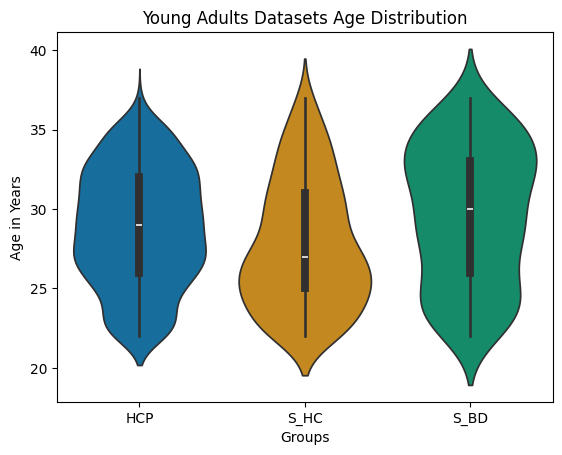 | 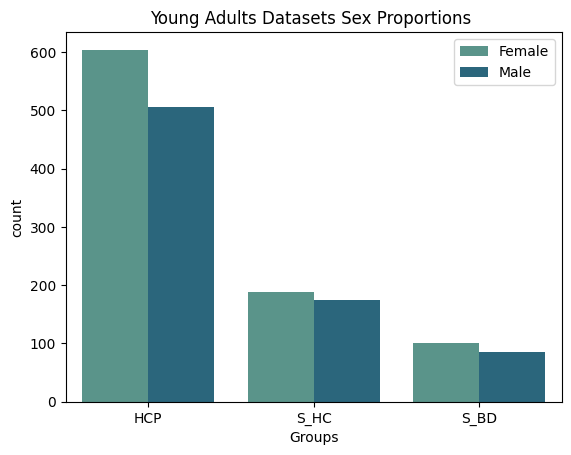 |
| --- | --- |

**Supplementary Figure 2 | Boxplot mean feature distributions across sites and groups****: pre vs. post harmonization**

Boxplots for mean CT, GMV, WMV, and TIV, displayed by site and by group, before and after data harmonization. The reference HCP site, id=0, remains unchanged while all the others are brought to its level (location and scale adjustment). The initial differences between HC and BD in the StratiBip test set are maintained after harmonization.

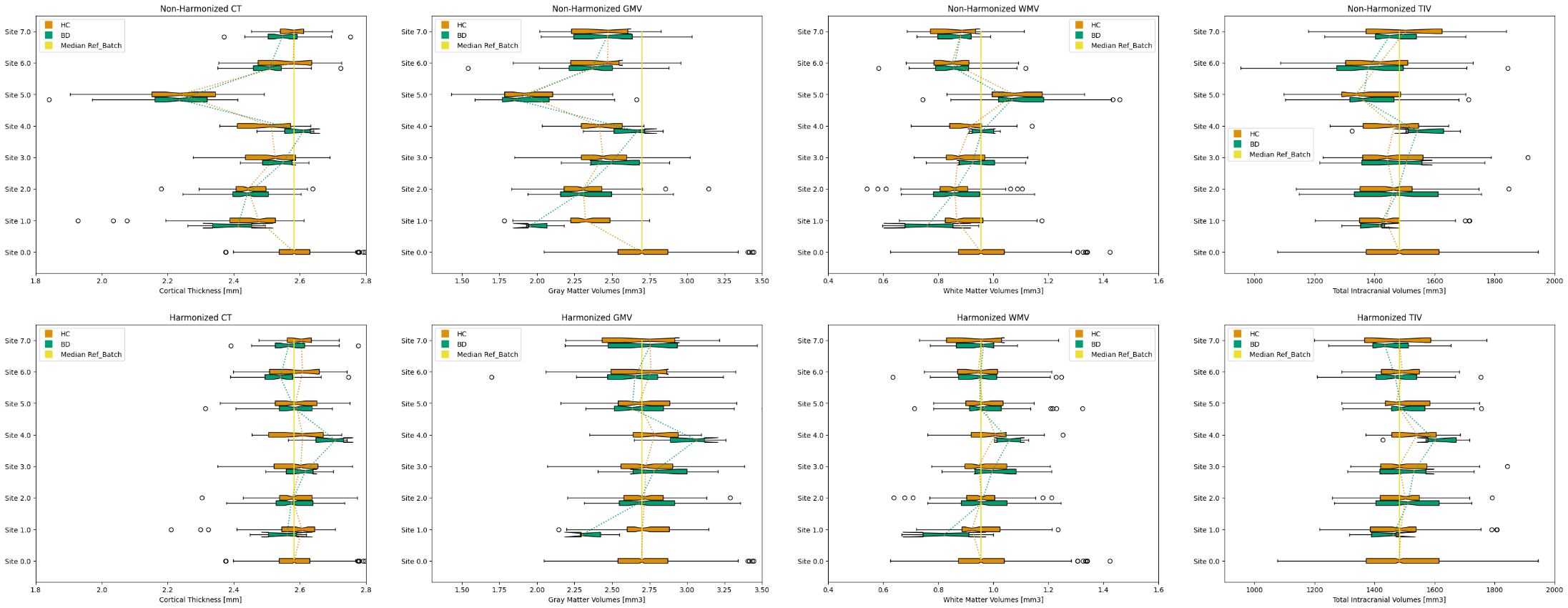

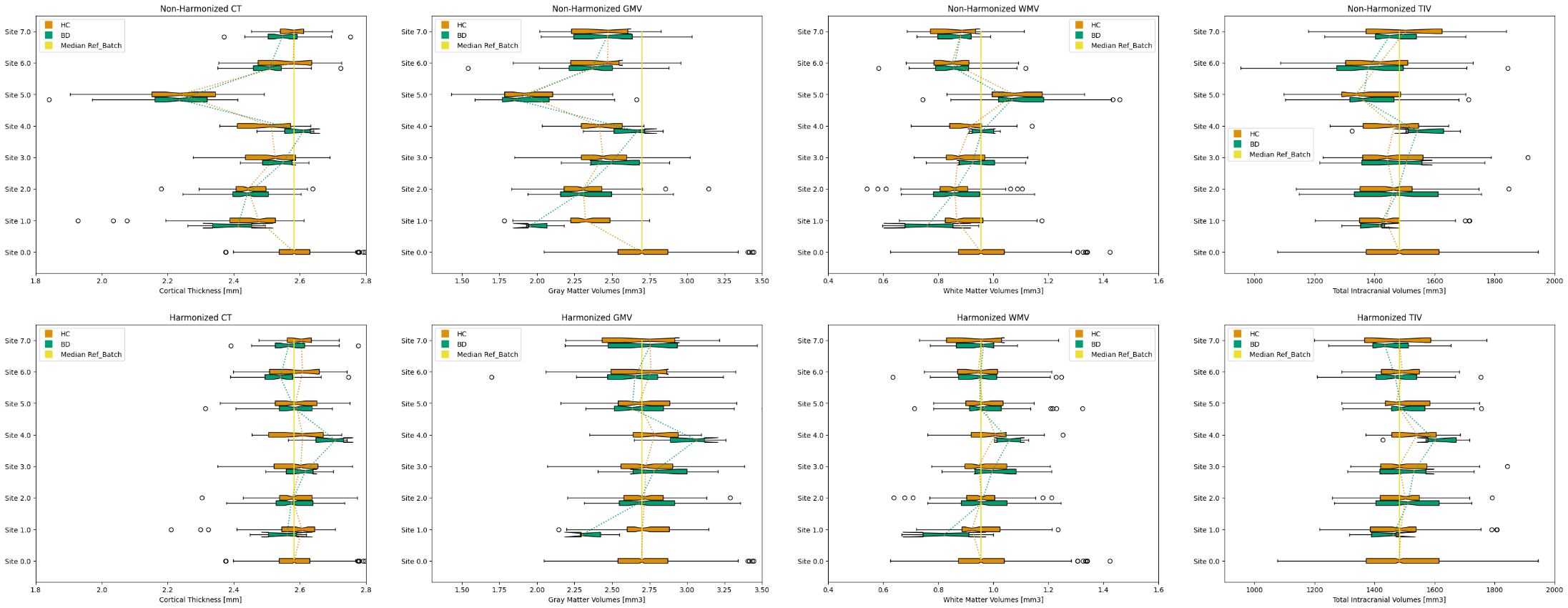

**Supplementary Figure 3 | Dataset UMAP 2D projections****: pre vs. post harmonization**

We assessed the UMAP 2D projections before and after harmonization, which revealed site-associated clusters before harmonization that effectively disappeared after harmonization. Top figure: projection of all dataset before (left) and after (right) M-ComBat harmoniation. Middle row figure: projection of globals data (TIV, absGMV, absWMV, average CT) for all sites, before (left) and after (right) harmonization. Two bottom figures: We inspected exclusively the BD-StratiBip data harmonization concatenated with the HCP, for both brain features and global features, and found the results to be equally satisfactory.

**Raw dataset	Harmonized dataset**

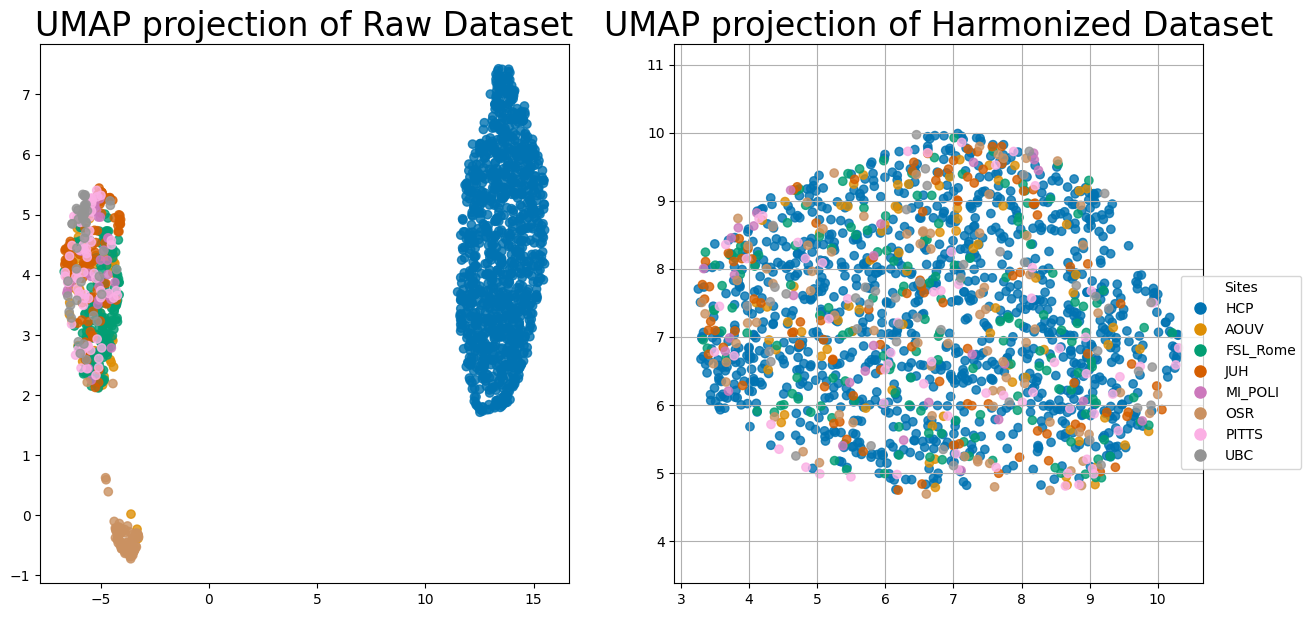

**Raw globals	Harmonized globals**

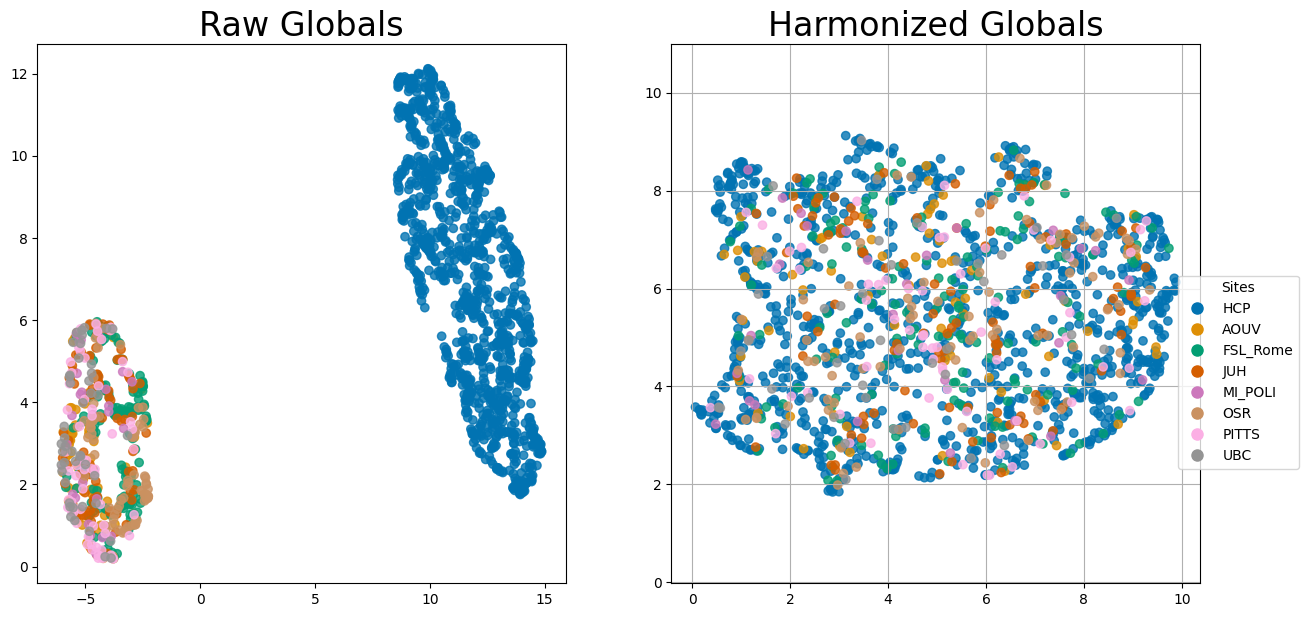

**Raw HCP + BD-StratiBip dataset	Harmonized HCP + BD-StratiBip**

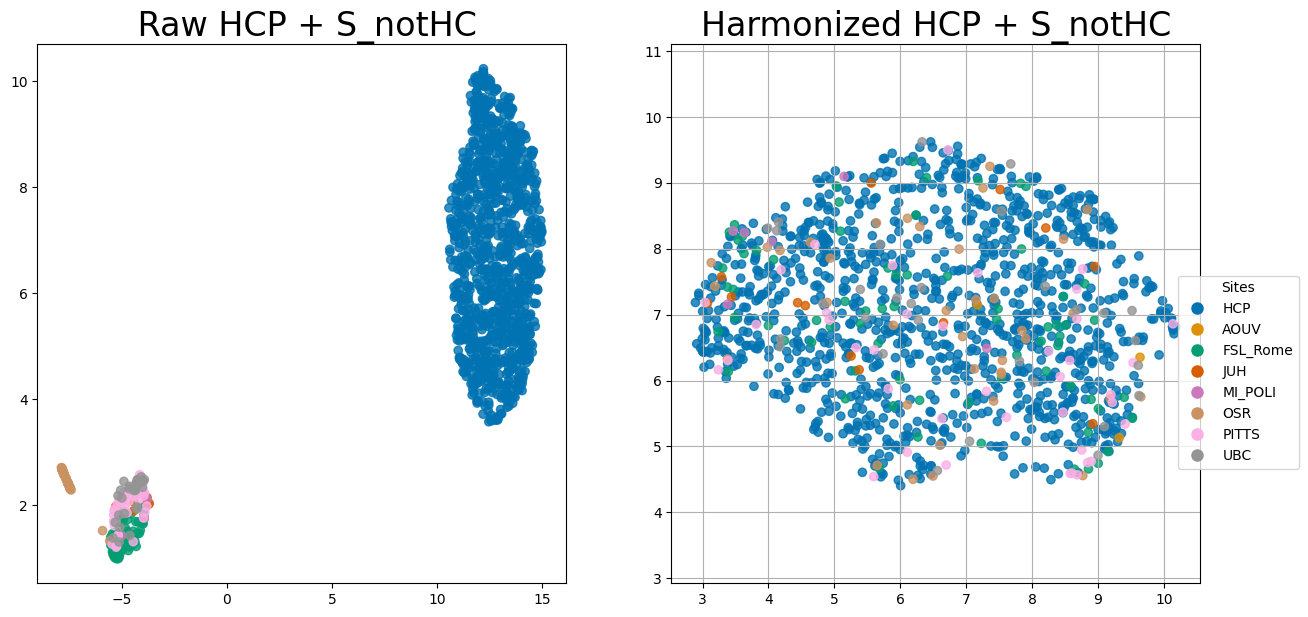

**Raw globals HCP + BD-StratiBip	Harmonized globals HCP + BD-StratiBip**

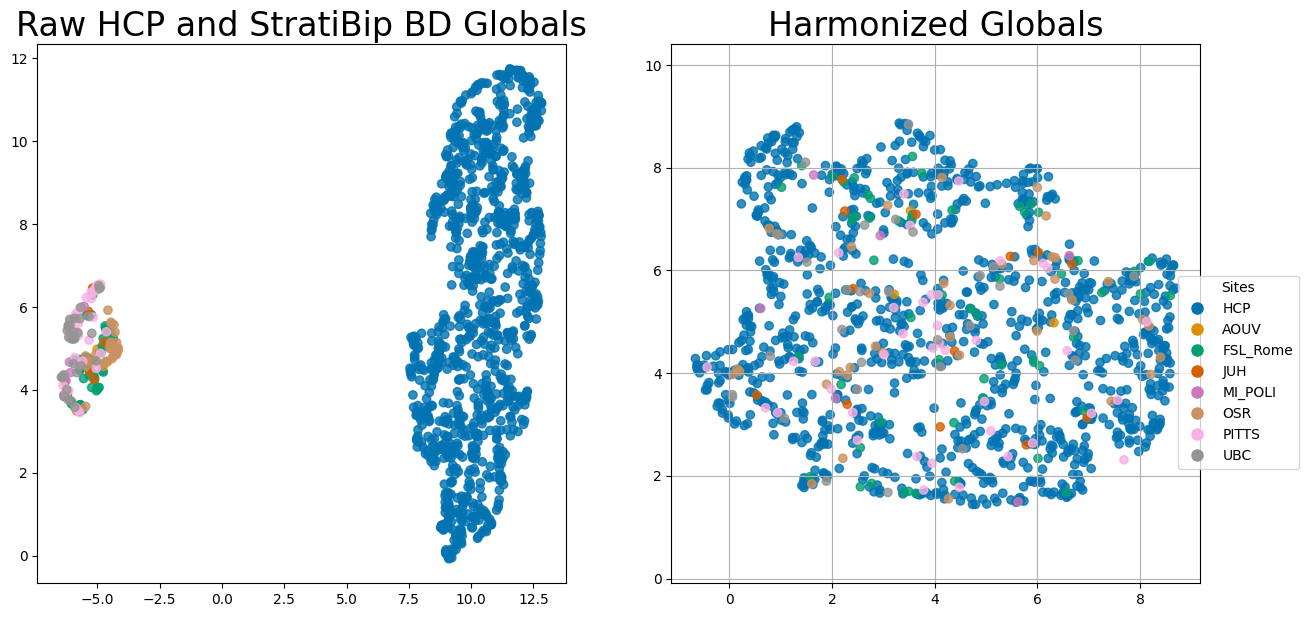

**Supplementary Figure 4 | AE model training evolution**

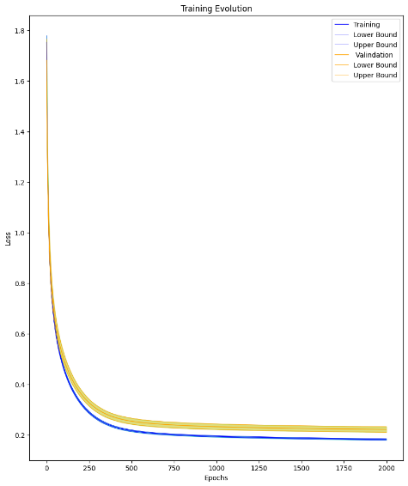

**Supplementary Fig.5 |** **95% CI Features MDS**

For each iteration of the bootstrap procedure, we calculated the feature-wise MDS based on the mean squared error (MSE), averaged by subject within HC or BD group. For each feature we report the MDS 95% CI.

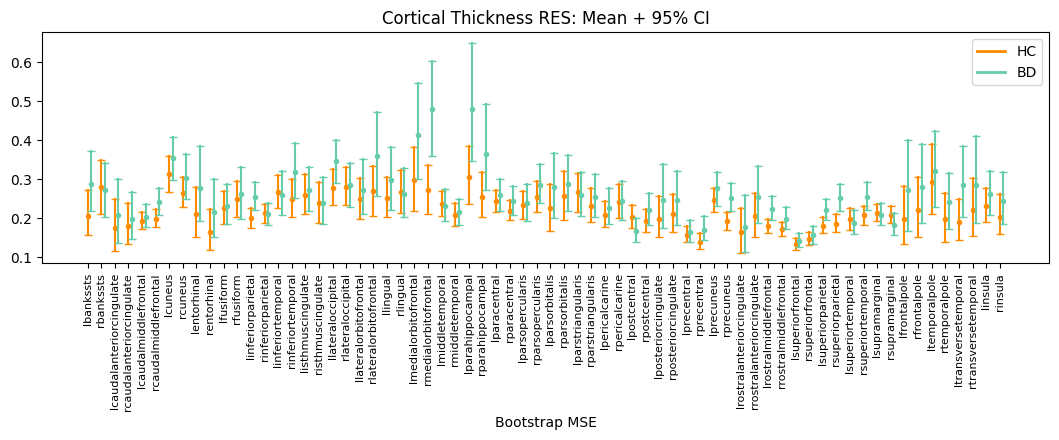

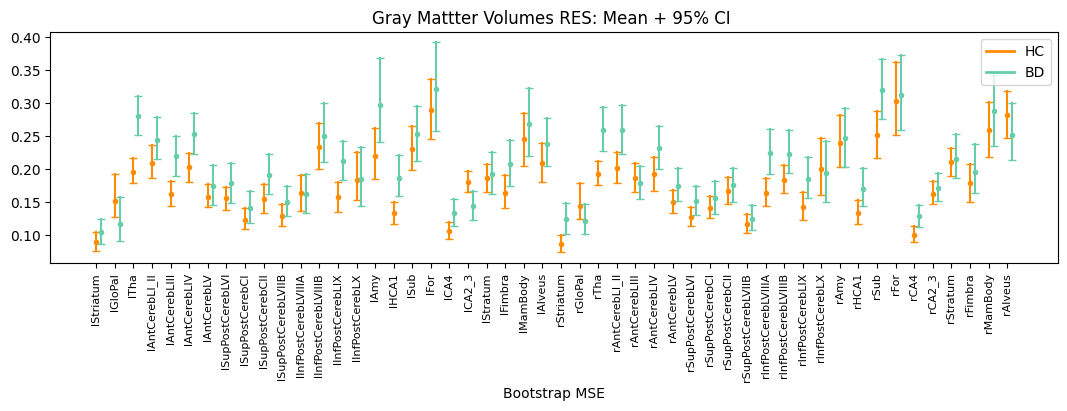

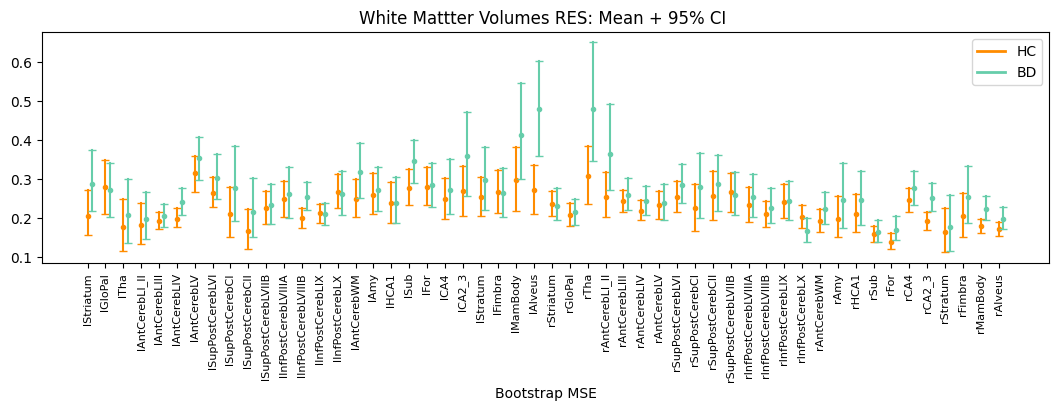

**Supplementary Fig. 6 | HCP-YA mZ scores features histograms with respective 99^th^ percentile threshold**

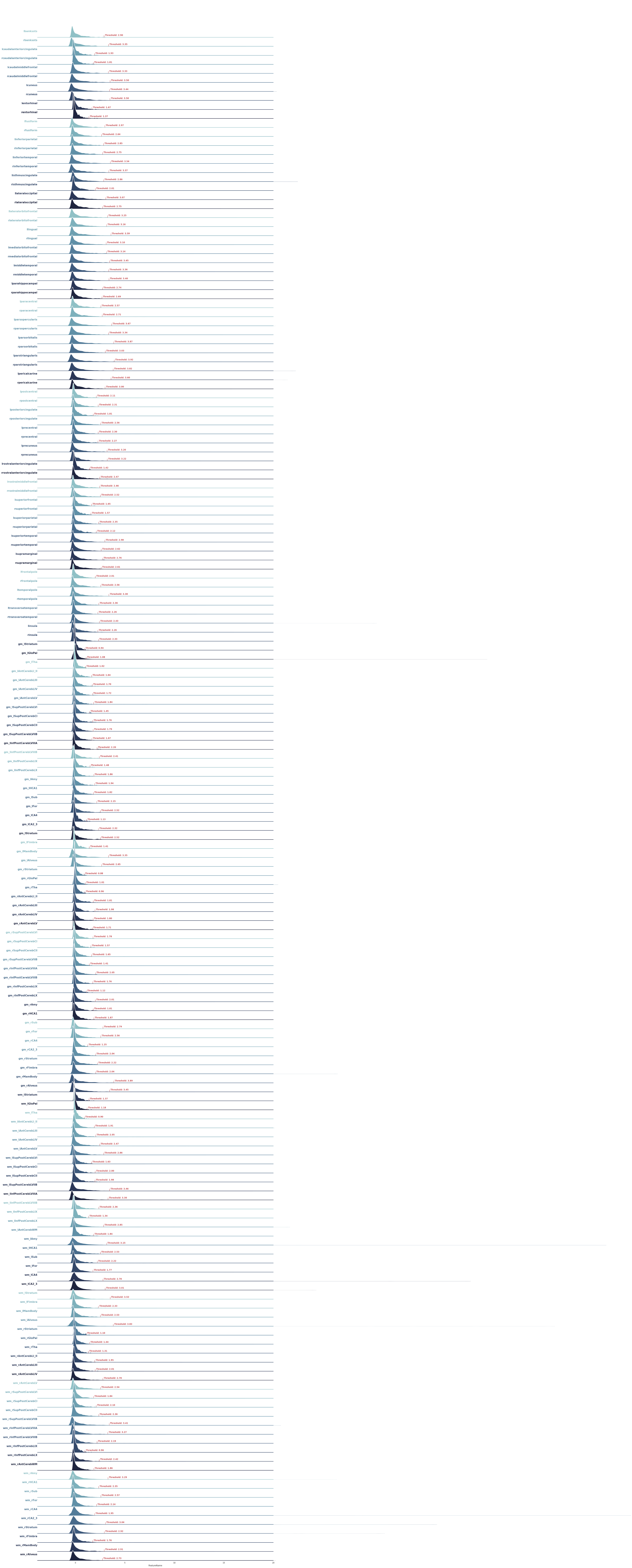

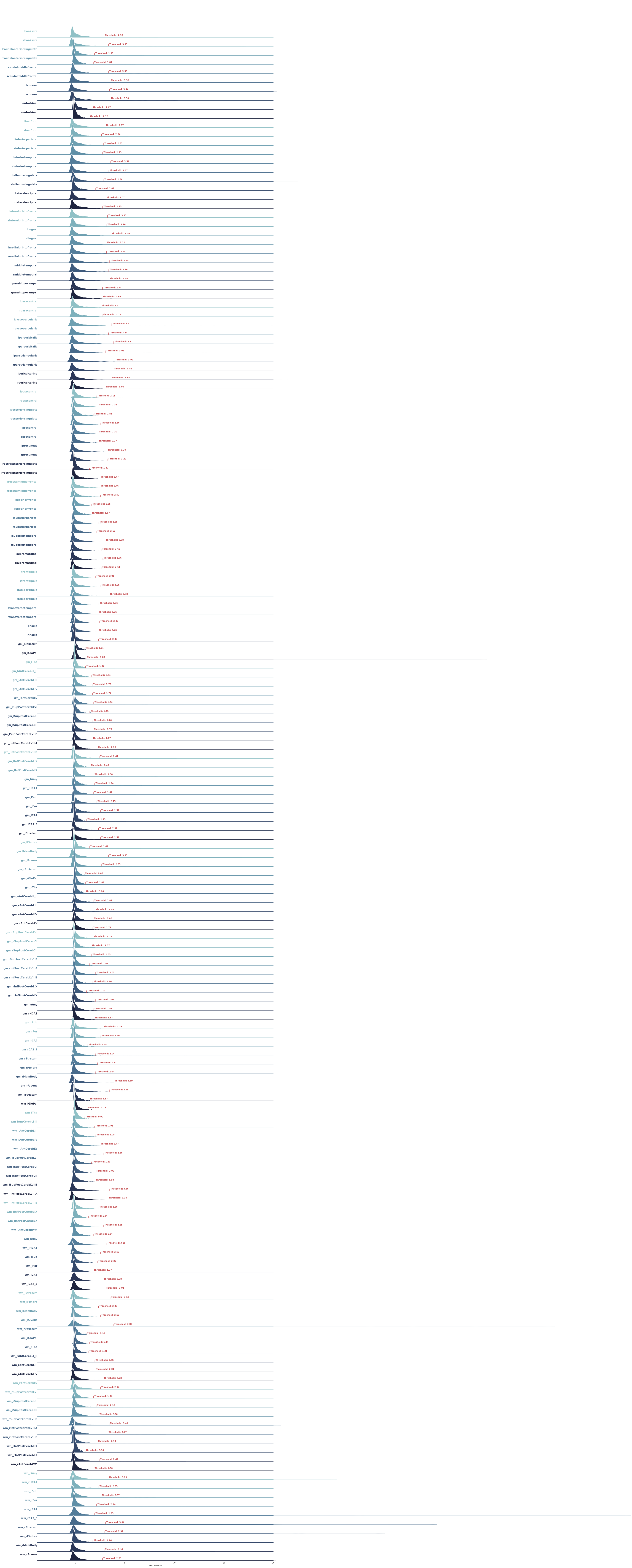

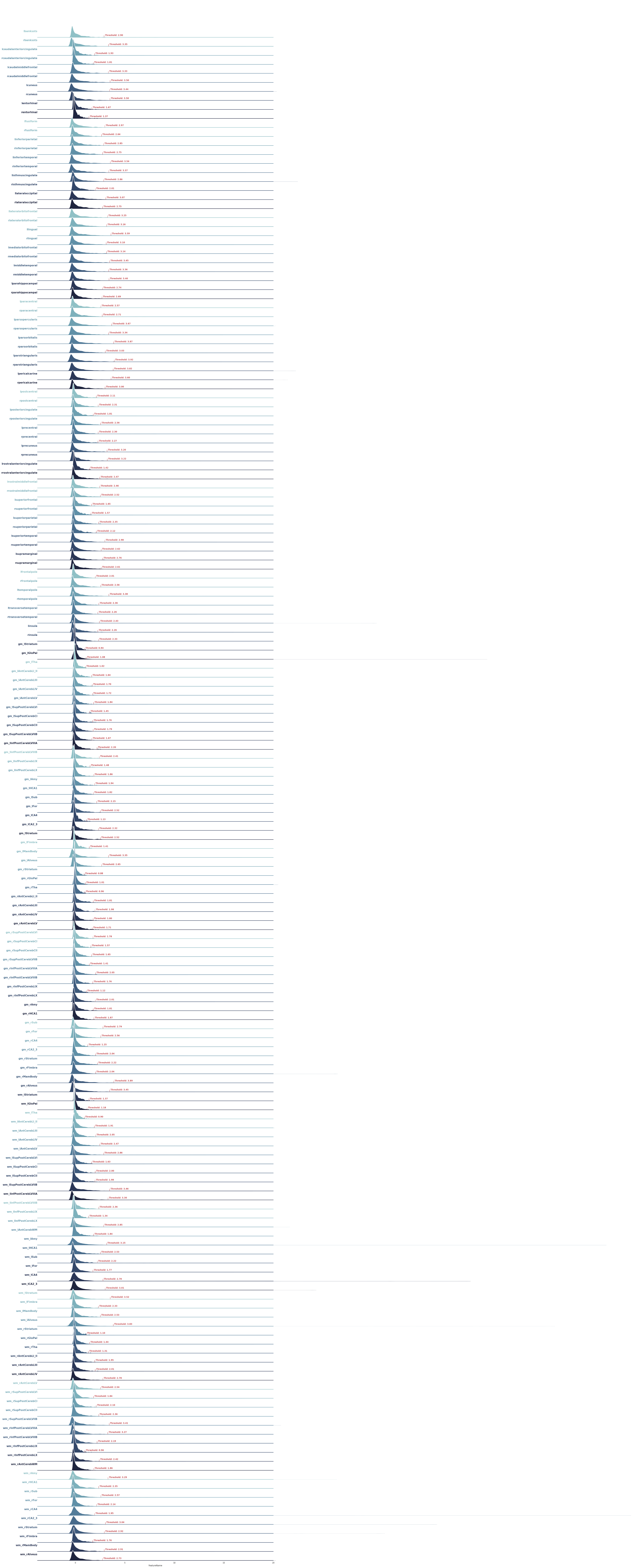

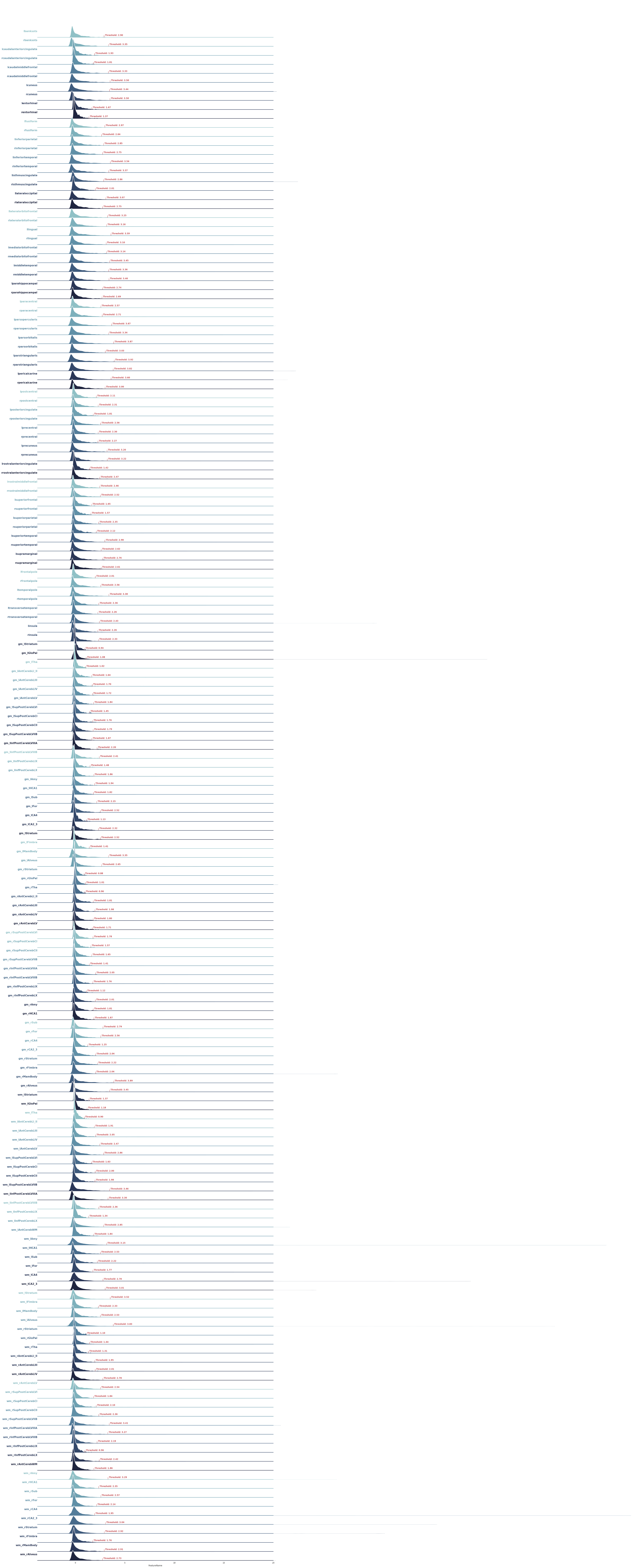

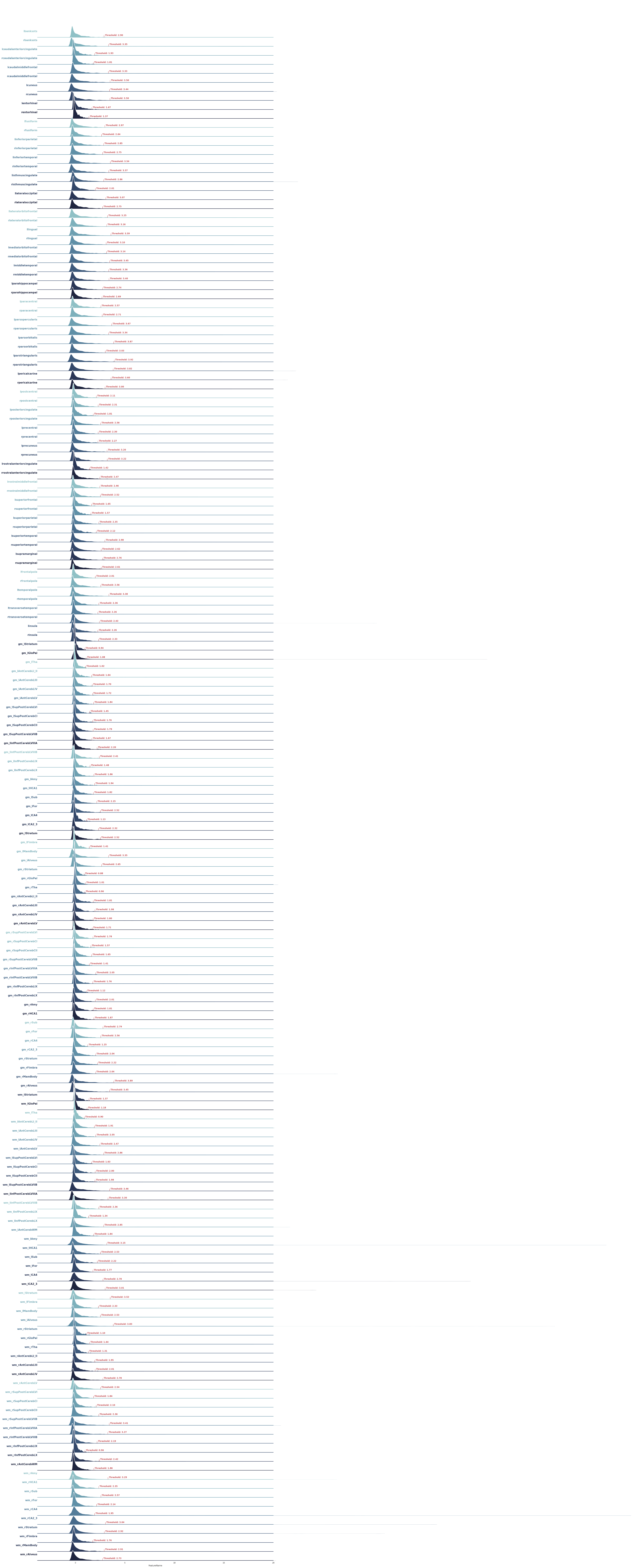

**Supplementary Fig. 7 | Features mZ distributions for HC and BD group**

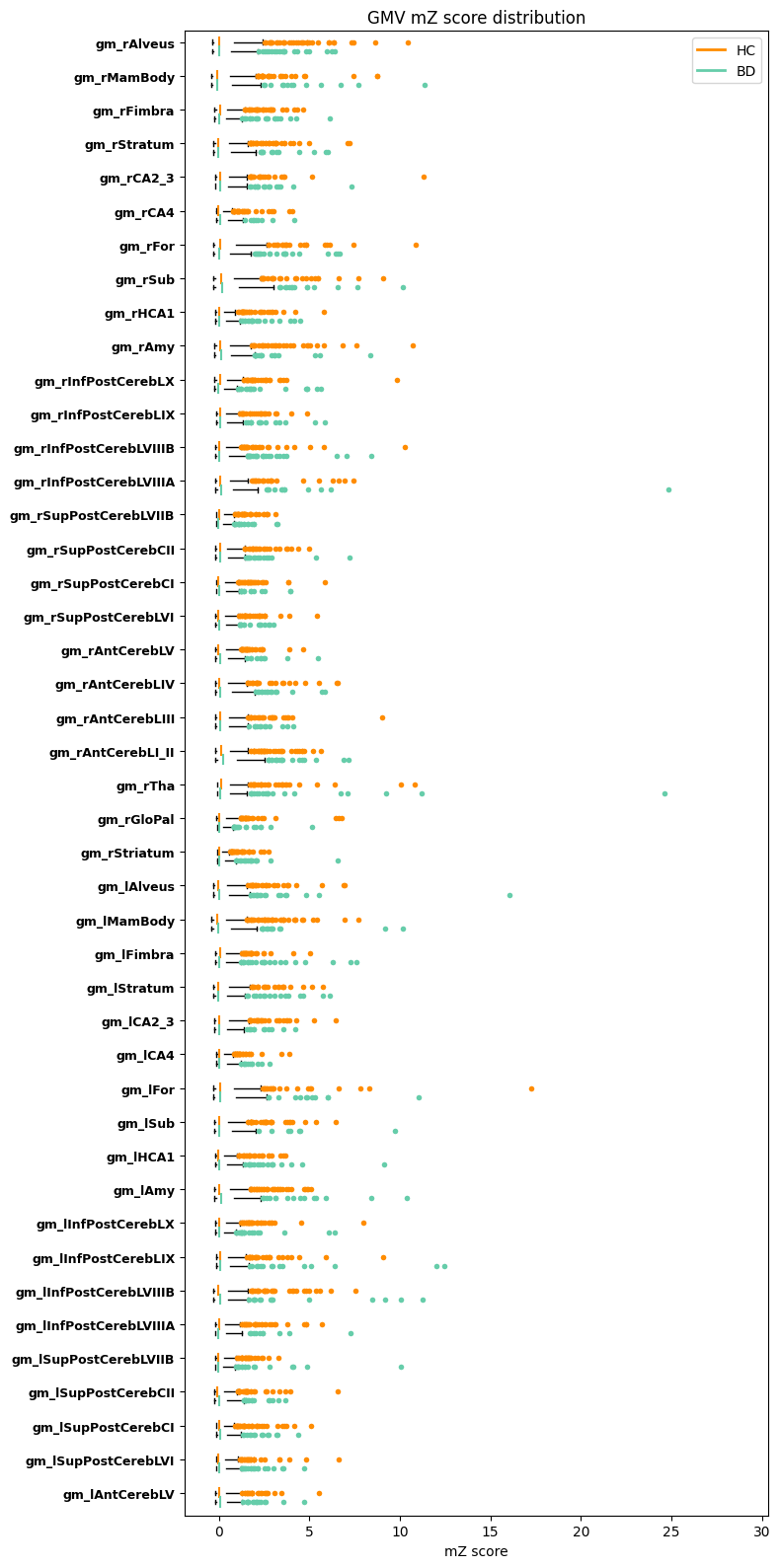

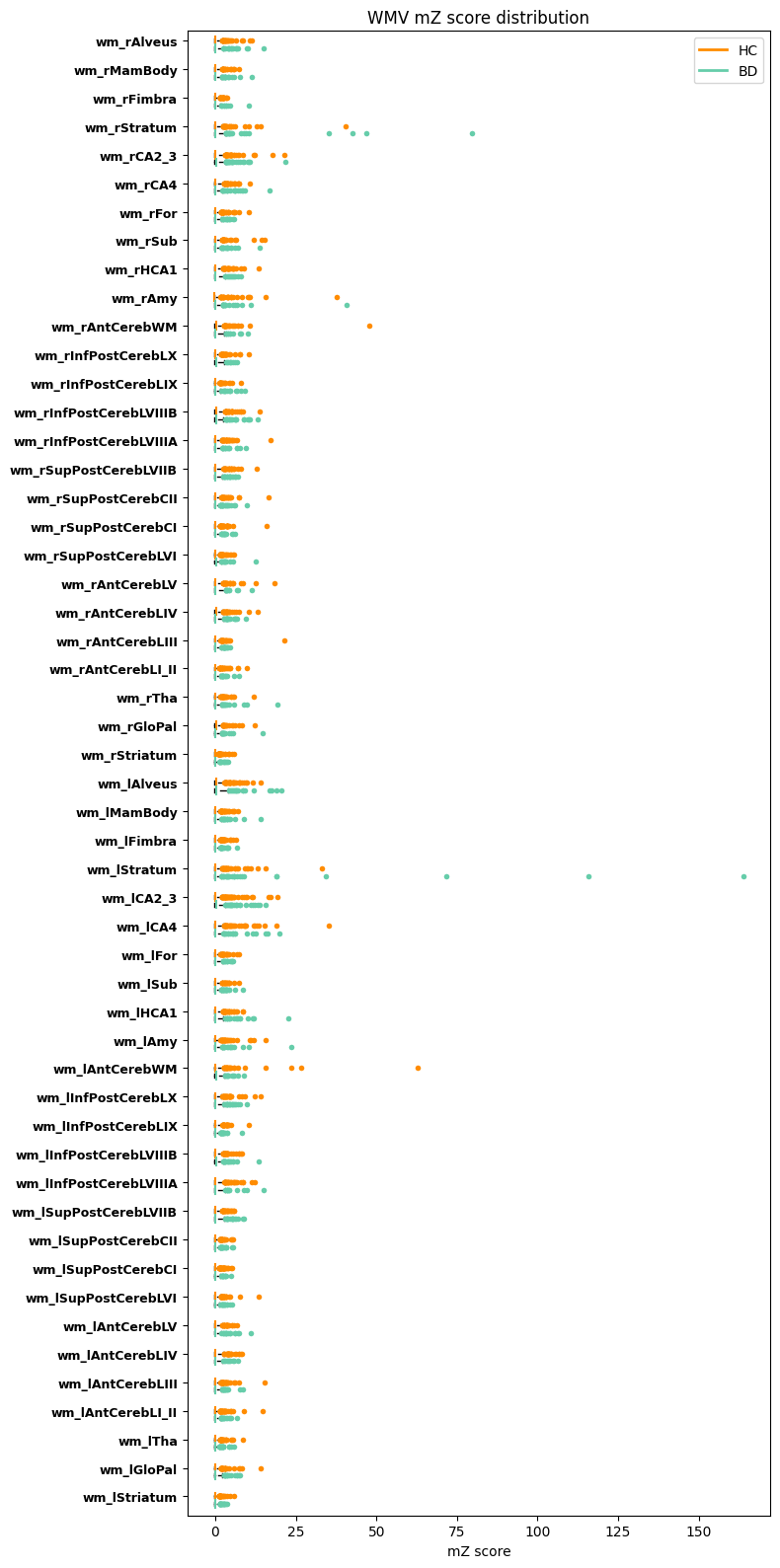

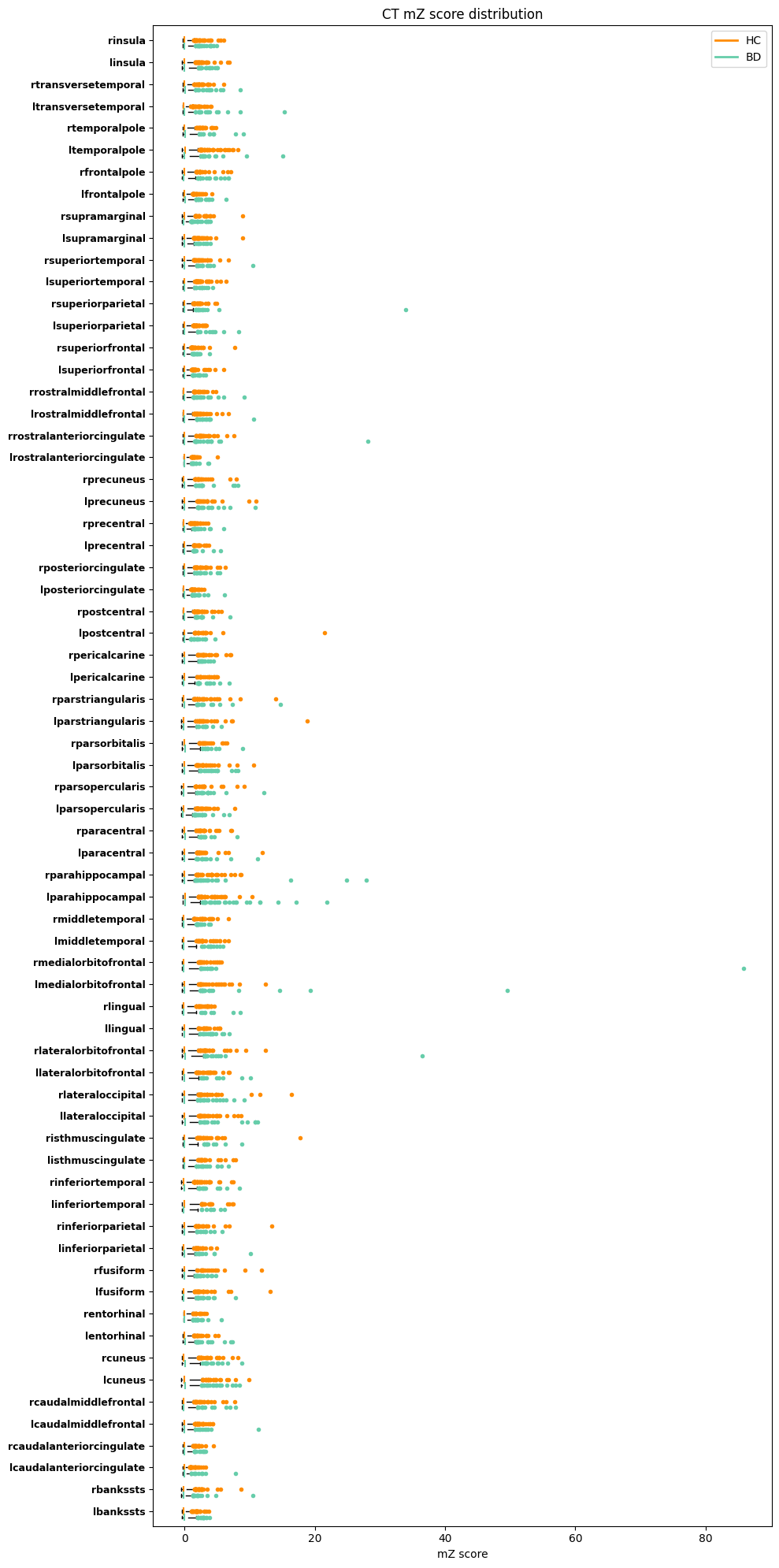
